# Alpha oscillations and stimulus-evoked activity dissociate metacognitive reports of attention, visibility and confidence in a rapid visual detection task

**DOI:** 10.1101/2021.11.23.469669

**Authors:** Matthew J Davidson, James S.P. Macdonald, Nick Yeung

## Abstract

Variability in the detection and discrimination of weak visual stimuli has been linked to oscillatory neural activity. In particular, the amplitude of activity in the alpha-band (8-12 Hz) has been shown to impact upon the objective likelihood of stimulus detection, as well as measures of subjective visibility, attention, and decision confidence. We aimed to clarify how preparatory alpha influences performance and phenomenology, by recording simultaneous subjective measures of attention and confidence (Experiment 1), or attention and visibility (Experiment 2) on a trial-by-trial basis in a visual detection task. Across both experiments, alpha amplitude was negatively and linearly correlated with the intensity of subjective attention. In contrast to this linear relationship, we observed a quadratic relationship between the strength of alpha oscillations and subjective ratings of confidence and visibility. We find that this same quadratic relationship links alpha amplitude to the strength of stimulus evoked responses. Visibility and confidence judgements corresponded to the strength of evoked responses, but confidence, uniquely, incorporated information about attentional state. As such, our findings reveal distinct psychological and neural correlates of metacognitive judgements of attentional state, stimulus visibility, and decision confidence.

## Introduction

This study explores the relationship between EEG alpha oscillations, objective performance, and subjective reports of visibility, attention, and confidence in a visual detection task, with two aims. The first is to characterise the relationship between preparatory alpha activity and objectively measured vs. subjectively experienced aspects of visual processing. Alpha (8-12 Hz) oscillations are prominent in spontaneous neural recordings, being readily observable to the naked eye (Berger, 1929). Rather than reflecting a passively idling state (Pfurtscheller et al., 1996), these oscillations are now recognised to account for a substantial portion of the behavioural variability that is recorded during psychophysical tasks (Ress et al., 2000). Recent M/EEG studies have shown that the power (Babiloni et al., 2006; Benwell et al., 2017; Ergenoglu et al., 2004; Iemi et al., 2017; Iemi & Busch, 2018; Limbach & Corballis, 2016) and phase (Busch et al., 2009; Coon et al., 2016; Mathewson et al., 2009; VanRullen et al., 2011) of prestimulus activity can determine perceptual outcomes. In related work, we have shown that alpha oscillations during active task preparation vary with task engagement as it fluctuates over time (Macdonald et al., 2011) and as a function of experimental manipulations such as reward (Hughes et al., 2013). Collectively, these results hint at the possibility of predicting perception and behaviour based on earlier neural states, although at present, the effects of alpha dynamics on objective and subjective measures of performance have been mixed. Here we aim to clarify how the strength of preparatory alpha band activity influence performance, subjective reports, and the generation of sensory evoked potentials (Chaumon & Busch, 2014; Hanslmayr et al., 2007; Hughes et al., 2013; Iemi et al., 2019; Min et al., 2007).

Our second, complementary aim is to use these neural markers—preparatory alpha and evoked potentials—to characterise the information that underpins subjective reports of attention and confidence. There is growing interest in the mechanisms and functional role of metacognitive processes that monitor and regulate ongoing processing (Fleming & Frith, 2014). Much of this work has focused on decision confidence—a subjective evaluation of the likelihood that a judgement reached is correct (Kepecs & Mainen, 2012; Yeung & Summerfield, 2012). According to influential theories, confidence reflects a readout of the strength of evidence in favour of the chosen option (Kepecs & Mainen, 2012; Pleskac & Busemeyer, 2010; Vickers & Packer, 1982). However, confidence is additionally sensitive to features such as the perceived reliability of evidence (Boldt et al., 2017), speed of decision (Kiani et al., 2014), and even social context (Bang et al., 2017), suggesting that evidence strength is combined with relevant contextual information in generating confidence reports (Shekhar & Rahnev, 2018). In parallel with this work on confidence, a separate body of research has investigated people’s introspective insight into their degree of attentional focus. Introspective reports of attentional state are predictive of objective performance across a range of tasks (Smallwood & Schooler, 2015), and correlate with neural markers including preparatory (Hughes et al., 2013; Macdonald et al., 2011) and prestimulus alpha (Whitmarsh et al., 2014, 2017, 2021; Worden et al., 2000), and stimulus-related potentials (Barron et al., 2011). Although some studies have begun to explore the relationship between attention and confidence (Denison et al., 2018; Kurtz et al., 2017; Rahnev et al., 2011; Recht et al., 2019, 2021; Zizlsperger et al., 2012), substantive questions remain, in particular regarding whether confidence reports incorporate contextual information about participants’ attentional state, and the degree to which subjective reports of confidence and attention depend on similar vs. distinct sources of information. We address these questions here.

Alpha oscillations provide an exciting opportunity to investigate relationships between attention, sensory processing and introspective reports. Recent studies suggest that alpha activity prior to the onset of a stimulus may govern objective performance criteria, albeit with somewhat inconsistent results. For example, higher prestimulus alpha power has been shown to either increase (Babiloni et al., 2006; Becker et al., 2011; Linkenkaer-Hansen et al., 2004), or decrease objective behavioural performance (Ergenoglu et al., 2004; Iemi et al., 2017; van Dijk et al., 2008), across a wide range of task contexts (Clayton et al., 2018; Iemi et al., 2017; Van Diepen et al., 2019). Similarly, visual detection has been shown to depend on when stimuli are presented relative to the phase of prestimulus alpha oscillations (Busch et al., 2009; Mathewson et al., 2009; VanRullen et al., 2011), perhaps dependent on the state of attentional focus (Kizuk & Mathewson, 2017; Mathewson et al., 2009). One partial account for these discrepancies, and a convergent theme within this literature, is that alpha oscillations reflect a state of relative cortical excitation or inhibition, which is mediated under top-down control to facilitate sensory processing (Van Diepen et al., 2019). In this context, weaker prestimulus alpha oscillations are indicative of a more highly excitable cortical state (Klimesch et al., 2007; Romei et al., 2008), which supports the negative relationship that has been reported between alpha amplitude and detection performance (Ergenoglu et al., 2004; Hanslmayr et al., 2005; van Dijk et al., 2008). Consistent with this view, prestimulus alpha oscillations are sensitive to attention, decreasing over cortical sites when attending to task-relevant information (Gould et al., 2011; Peylo et al., 2021; Sauseng et al., 2005; Thut et al., 2006).

More recently, however, evidence has linked alpha oscillations to the subjective aspects of visual decisions, which may bias behavioural performance in lieu of any change in sensory precision (Benwell et al., 2017; Limbach & Corballis, 2016; Samaha, LaRocque, et al., 2020). In particular, low prestimulus alpha power has been shown to precede a higher incidence of target detection and false-alarms (Iemi et al., 2017; Limbach & Corballis, 2016; Samaha, Iemi, et al., 2020) suggesting that low alpha power may improve detection performance only indirectly, by biasing participants to report ‘yes’ in a detection task regardless of the veridical presence of a target stimulus. In support of this view, in two recent examples, the strength of alpha power preceding a two alternative forced choice (2AFC) discrimination task was shown to negatively correlate with decision confidence (Samaha et al., 2017), and perceptual awareness/target visibility (Benwell et al., 2017) without any change in objective accuracy.

Variations in the strength of alpha oscillations have thus been associated with changes in objectively measured and subjectively reported indices of sensory processes and attention. To decouple the influence of alpha on these overlapping indices, we analysed data from two EEG experiments involving a near-threshold target detection task, in which we collected simultaneous ratings of both decision confidence and attention (Experiment 1), and target visibility and attention (Experiment 2) on a trial-by-trial basis. Our analysis focused on how preparatory alpha amplitude impacts upon these outcomes during a rapid target detection task. For both experiments we used an identical stimulus detection task involving decisions about stimulus presence/absence - decisions that have distinct neural contributions (Mazor et al., 2020), and metacognitive correlates (Kanai et al., 2010; Meuwese et al., 2014) compared to their 2AFC counterparts. Combined, the results of the two experiments allow us to assess how preparatory alpha activity influences sensory processing and introspective reports. Contrasted, the results of the two experiments provide insights into the contribution of attention and sensory evidence to judgements of confidence (Experiment 1) and stimulus visibility (Experiment 2).

To preview our results, we show that participants’ confidence reports (but not their ratings of stimulus visibility) correlate with their self-reported attentional state, suggesting a partial dependence of the two key forms of introspective report. This correlation notwithstanding, our EEG analyses indicate that evaluations of confidence and attention depend on partially distinct sources of information: We demonstrate that a quadratic, inverted-U function links preparatory alpha amplitude to subjective visibility and confidence in a detection task, whereas attention negatively and linearly correlated with alpha amplitude. We further show that both confidence and visibility increase with the strength of visually evoked potentials, which were also quadratically modulated by preparatory alpha amplitude.

## Materials and Methods

### Participants

A total of 21 participants participated in this research, 12 participants in Experiment 1, and 9 in Experiment 2. A portion of the data from Experiment 1 has previously been published (Macdonald et al., 2011). That work showed that single-trial ratings of subjective attention could be classified based on preparatory alpha power, and that this classification was optimal over a sliding-window of several minutes (Macdonald et al., 2011). Here, we focus instead on how preparatory alpha amplitude (and, in Supplemental analyses, the phase of this activity) affect the generation of target-evoked event related-potentials (ERPs), and the interaction of preparatory alpha and ERPs on subjective criteria. Experiment 2 is a new experiment. There were five males in Experiment 1, and all participants’ ages ranged from 18-29 years (*M* = 22.3, *SD* = 4.4). There were 4 males in Experiment 2, and all participants’ ages ranged from 19-23 years (*M* = 20.6, *SD* = 1.8). All participants were recruited for participation at the University of Oxford, were paid for their participation, and had normal or corrected to normal vision. This research was conducted in accordance with the University of Oxford’s institutional review board, and the American Psychological Association’s standards for ethical treatment of participants.

### Experimental procedure

The experimental procedure was very similar between the two experiments, and has previously been detailed in Macdonald et al., (2011). In each trial, participants were asked to monitor a rapid serial visual presentation (RSVP) of images for a difficult-to-detect target image. Each trial began with the words ‘Get Ready’ presented on screen for 300 ms, before the 10 images comprising the RSVP stream were presented after a further 700 ms. Each image in the stream was presented for 50 ms, followed by a blank interval for 50 ms, resulting in a 10 Hz presentation rate. Each image was a grey-scale pattern of white noise, and target images included a set of six superimposed concentric circles (each subtending 0.4° visual angle), arranged in a hexagonal pattern (subtending 3.3° visual angle; Figure 1). There were 936 trials in total. Targets were presented on 50% of trials, with their position in the RSVP stream balanced across image positions 3-8. For each participant, the contrast of the hexagonal target pattern was determined in a pre-experimental session to titrate detection rates to approximately 75% (QUEST, Psychophysics Toolbox 3, (Brainard, 1997)).

**Figure 1.**
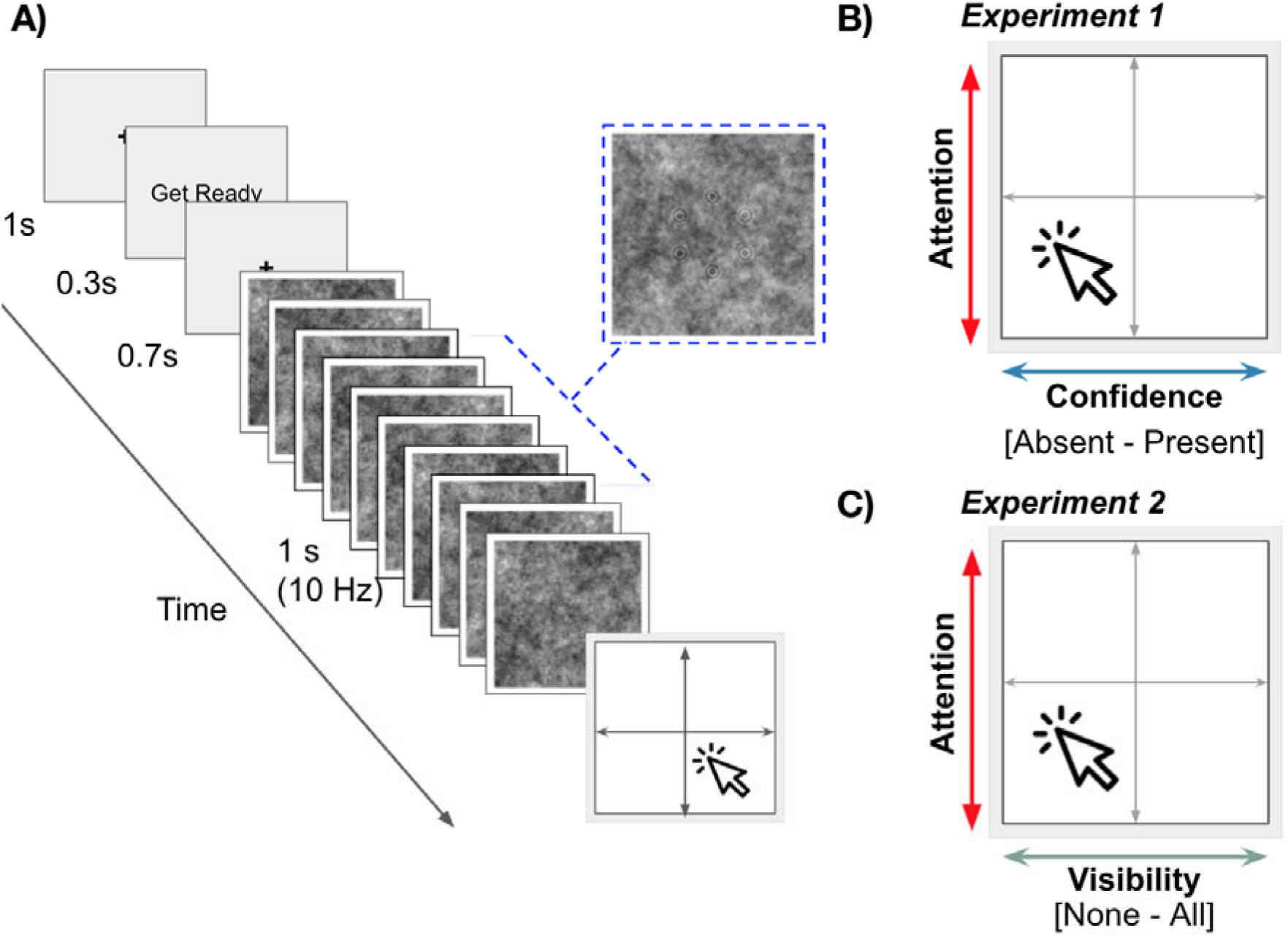
Trial procedure and response options. A) Each trial began with the words ‘Get Ready’ presented on screen. After a fixed interval of 1s, the RSVP sequence began, and a target image was presented once, on 50% of trials. Targets (shown outlined in blue) were presented in one of positions 3 to 8 in the RSVP stream. B) After each trial, participants rated either their subjective confidence and attention (Experiment 1), or C) the perceived visibility of the target and their attention (Experiment 2).

After the RSVP stream, participants indicated their subjective attention and confidence (Experiment 1), or attention and visibility (Experiment 2) ratings by providing a single mouse-click within the response screen (Figure 1). In Experiment 1, the response screen was subdivided into four quadrants by faint grey lines, with the prompts “Did you see the target?”, “How confident are you of that?” and “How focused were you?” Displayed at the top of the screen. The words “Sure Absent” and “Sure Present” were presented on the left and right extrema of the x-axis, and “More Focused” and “Less Focused” placed on the top and bottom of the y-axis. In Experiment 2, the prompt at the top of the screen replaced the question about confidence with one targeting stimulus visibility: “How much of the target did you see?”, with extremes of the x-axis labelled as “None” and “All”. In both experiments, the response screen was 201 x 201 pixels. Attention was measured on a 201-point scale according to the y-axis click location. In Experiment 1, confidence in presence or absence was measured on a 100-point scale (decreasing or increasing distance from the vertical midline), and in Experiment 2 visibility was measured on a 201-point scale according to the x-axis click location.

Participants were instructed to rate their subjective state only with respect to the current trial, and to incorporate their attention and confidence/visibility in this single response. Thus, in Experiment 1, the horizontal distance from the vertical midline represents confidence in the presence or absence of a target, and in Experiment 2 click distance from the left extrema represents target visibility. In both experiments, click position on the vertical axis represents trial-specific attention to the detection task.

### Behavioural analysis

For our behavioural analysis, we calculated overall target accuracy (% correct), as well as hit rate (HR; “Yes” responses in target-present trials), false alarm rate (FAR; “Yes” responses in target-absent trials), and standard metrics from signal detection theory (Green & Swets, 1966). Hits and false alarms (i.e., trials on which participants’ responses were taken to indicate a target was present rather than absent) were defined in Experiment 1 as clicks in the right half of the response screen, and in Experiment 2 as any click away from the left extrema of this screen. We calculated d’, which measures the sensitivity between signal and noise distributions in the signal detection framework, as well as decision criterion (*c*), which measures the likelihood of “Yes” responses, regardless of the veridical presence of a stimulus. When *c* is positive, the decision criterion is said to be conservative, and negative *c* values indicate a more liberal criterion - or tendency to respond ‘yes’ in detection tasks, relative to the true unbiased response probability given by the intersection between signal and noise distributions.

We also calculated metacognitive sensitivity (type-2 performance), which captures the fidelity of introspective judgements with relevance to objective performance. High type-2 performance indicates that introspective judgements are well-calibrated, and positively correlated with the objective likelihood of a correct response. Low type-2 performance indicates that introspective judgements are a poor indicator of objective accuracy. We quantified type-2 performance as the area under the ROC curve (AUROC2; (Fleming & Lau, 2014)), constructed from each participant’s subjective confidence, visibility, or attention ratings: Specifically, for every rating value used by a particular participant, we calculated the proportion of all correct response trials and the proportion of all incorrect response trials with ratings that exceeded this value, and then calculated the area under the curve created by plotting these proportions (on the y- and x-axis, respectively) for all rating values. A value of 1 indicates perfect sensitivity; a value of 0.5 indicates chance performance.

### EEG recording and preprocessing

EEG was recorded from 32 Ag/AgCl electrodes using a Neuroscan Synamps 2 system. Electrode positions were FP1, FPz, FP1, F7, F3, Fz, F4, F8, FT7, FC3, FCz, FC4, FT8, T7, C3, Cz, C4, T8, TP7, CP3, CPz, CP4, TP8, P7, P3, Pz, P4, P8, POz, Oz, Oz, and O2. During recording, all electrode impedances were kept below 50 kΩ. Four additional electrodes were placed over the outer canthi of left and right eyes, and above and below the right eye to measure eye-movements. Two additional electrodes were attached to the left- and right mastoids, of which the left acted as a reference. All EEG data were recorded at a sampling rate of 1000 Hz, before being downsampled off-line to 250 Hz, and low pass-filtered at 48 Hz. EEG data were epoched from 0.5 s before, to 3 s after the onset of the words ‘Get Ready’ on screen and demeaned using the whole-epoch average. Noisy channels were identified by visual inspection and replaced with the average of nearest neighbours. In Experiment 1, an average of 0.25 channels were removed (3 over all participants), and no channels were removed in Experiment 2. Independent component analysis was performed to identify and remove artefacts using the SASICA toolbox (Chaumon et al., 2015), and all epochs were visually inspected for rejection. On average <4% of trials were discarded per participant.

### Preparatory Alpha Analysis

Analysis was performed within MATLAB (R2019a) using custom scripts, and functions from the EEGlab (Delorme & Makeig, 2004), FieldTrip (Oostenveld et al., 2011), and Chronux (Bokil et al., 2010) toolboxes. Our analysis focused on alpha activity in the preparatory window, covering 1 s between the presentation of the words ‘Get Ready’ and onset of the RSVP stream, as well as the amplitude of ERPs evoked by the RSVP stream. Alpha oscillations, measured over 1 s between the words ‘Get ready’ and the onset of the RSVP stream, were strongest over parieto-occipital electrodes (POz, O1, Oz, O2). We averaged over these electrodes for all our alpha analyses.

To avoid the possibility of post-stimulus activity (i.e. the RSVP response) contaminating our measure of alpha band activity within the preparatory window, we avoided the use of a sliding window spectrogram (e.g. (Davidson et al., 2020)), or time-frequency decomposition via wavelet transform (e.g. (Benwell et al., 2017; Iemi et al., 2017)). Instead, single-trial alpha amplitude was calculated by applying the Fast Fourier Transform (FFT) to the Hanning tapered preparatory period in each epoch. We used a single taper per frequency (zero padded, resolution: 0.24 Hz), and retained the complex values of the FFT. We quantified the strength of alpha band activity by taking the absolute of these complex values, and estimated preparatory alpha amplitude by averaging these values over 8-12 Hz at each channel. Across both experiments, all participants’ peak alpha frequency fell within this canonical band (Experiment 1, *M* = 10.53 Hz, *SD* = 1.06; Experiment 2; *M* = 10.06 Hz, *SD* = 0.76). To facilitate comparisons across participants, we first applied the z-transform to all single-trial estimates of alpha amplitude per participant. We sorted single-trial values of alpha amplitude into quintiles, by binning according to the 0-20%,21-40%, 41-60%, 61-80% and >80% values of the cumulative probability distribution of z-transformed data. When sorting by a subclass of outcome (e.g., Hits only), we applied the quintile split after first restricting to the range of relevant trials, to ensure approximately equal trial numbers in each quintile bin. We performed the same quintile separation and binning procedure when also analysing behavioural and ERP responses by subjective criteria. When visualizing the power spectrum across frequencies (Figure 4), we additionally squared the complex values of our FFT and applied the log transform.

### ERP analysis

After sorting trials according to quintiles of alpha amplitude per participant, we next characterised how preparatory alpha modulates the event-related potentials evoked by the RSVP stream. Based on previous research, we focused on two measures: the early sensory-evoked P1 component elicited by the first image of each RSVP stream (which never contained a target stimulus) and the centro-parietal positivity (CPP or P300) elicited by detected targets. We use the P1 as a measure of the overall excitability of sensory cortex, and evaluate the CPP to detected targets as a measure of the strength of evidence associated with those targets (Murphy et al., 2015; O’Connell et al., 2012; Twomey et al., 2015).

We closely followed the analysis procedures detailed by (Rajagovindan & Ding, 2011), to investigate whether alpha affected early stimulus processing. To quantify the amplitude of the P1 component, each whole-trial preprocessed epoch was additionally filtered between 1 and 25 Hz (one-pass zero phase, hamming-windowed FIR filters), and a pre-RSVP baseline correction was applied using the period −50 to 0 ms relative to RSVP onset (950 to 1000 ms relative to the start of each trial). P1 amplitude was calculated by first averaging all trials within each alpha quintile, and then retaining the maximum positive peak within the window 80 to 160 ms after RSVP onset. We observed a reliable P1 component (i.e., positivity in 80-160 ms, across all 5 quintiles), only at the most occipital electrode sites (O1, Oz, O2), and report the average P1 amplitude averaged across these electrodes.

We also averaged the ERP response to targets which were embedded within the RSVP stream on target-present trials, focusing on the CPP that is thought to reflect the accumulating evidence for a decision. Target-locked ERPs were calculated after filtering preprocessed epochs between 0.1 and 8 Hz to remove the influence of the 10 Hz RSVP component (one-pass, zero phase, hamming-windowed FIR filters). We then sub-selected the period −200 ms to 1.5 s relative to target onset, and baseline corrected using the −100 ms to target onset window. When targets were presented within the RSVP stream (“Hits” and “Misses”), we quantified the CPP strength by averaging over a cluster of centro-parietal electrodes (C3, Cz, C4, CP3, CPz, CP4), over the period 250-550 ms relative to target onset.

### Mixed-effects analyses

One of our key motivations was to assess the effect that the strength of preparatory alpha band activity, split into quintiles, had on subjective measures. Because we observed a mixture of both linear and quadratic trends, we utilised mixed-effects models to formally test the nature of these trends, in preference to other analysis options such as a repeated-measures ANOVA. Our justification for this choice is two-fold. First, mixed effects models allow us to account for variance which is attributable to either individual participants (random effects), or a relevant category (e.g., fixed effects of alpha). Second, and most importantly, mixed-effect models are more appropriate to our research question, as by specifically testing either a linear or quadratic model, we can explicitly compare which may be a better fit to the data.

We formally tested the nature of linear and quadratic coefficients by performing a series of stepwise mixed effects analyses to model either linear or quadratic fixed effects of alpha amplitude, which included random effects (intercepts) per participant. We performed likelihood ratio tests between the full model, which combined random, linear, and quadratic effects, to restricted models of increasing simplicity (removing first the quadratic, and then linear term). We compared the goodness-of-fit for each model using likelihood ratio tests, and in our results report when either the linear or quadratic model was a better fit to the data than the basic model, which included only random effects per participant. When a significant linear or quadratic effect is reported, the fixed effect coefficient (β) and 95% confidence intervals are also included.

## Results

We recorded continuous measures of both confidence and attention (Experiment 1), and visibility and attention (Experiment 2) on a trial-by-trial basis in a visual detection task. We first present the behavioural results from these tasks, showing an asymmetry in the behavioural correlations between subjective measures and objective performance. We then report how differences in these performance measures are influenced by preparatory alpha amplitude. Finally, we show that alpha amplitude quadratically modulates the generation of sensory-evoked potentials, which in turn correlates with confidence and visibility judgements.

### Behavioural Results

#### Confidence and visibility correlate differently with attention ratings

The use of subjective responses and introspective accuracy varied between Experiments 1 and 2. After each trial, participants were asked to indicate either their trial-specific confidence and attention ratings (Experiment 1), or visibility and attention ratings (Experiment 2) by providing a single mouse-click within a response square. Figure 2A displays the cumulative total click responses in both experiments. Trials in which targets were presented within the RSVP stream are shown in orange, trials without a target are shown in purple. Figure 2 plots data pooled across all participants, but key trends apparent here are mirrored in single-participant data (see Supplementary Figures 1 and 2), despite typically-observed idiosyncratic differences across participants in their use of subjective rating scales (cf. (Ais et al., 2016).

**Figure 2.**
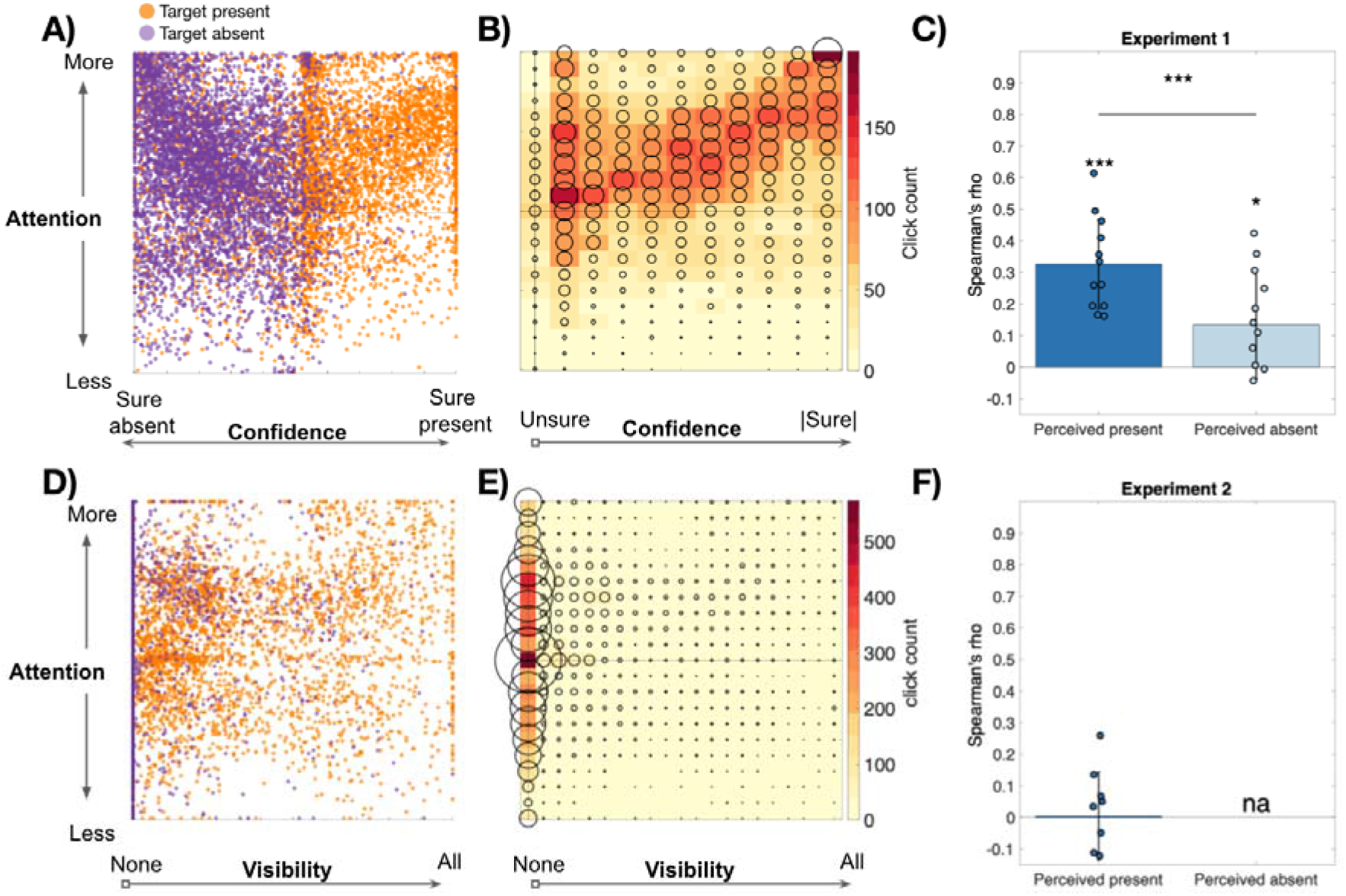
Subjective responses to the same visual detection task. **A)** In Experiment 1, participants rated their decision confidence that a target was either absent or present, simultaneously with their subjective attention, with a single click in the response square. Orange dots indicate target present trials, purple dots represent target absent trials. **B)** Increases in subjective confidence positively correlated with an increase in attention. **C)** Average linear correlation coefficients were significantly positive for attention and perceived-presence (orange), as well as attention and perceived-absence (purple). Error bars display 1 SD. **D)** In Experiment 2, participants rated the subjective visibility of targets on the x-axis. Colour conventions are the same as in **A-C). E-F)** Subjective visibility did not positively correlate with attention ratings. Note: no correlation is calculated for “Perceived absent” trials in Experiment 2 because these trials were defined as having the same (zero) visibility rating on all trials.

In Experiment 1, participants rated their confidence in the presence or absence of a target on the x-axis, and attentional state rating on the y-axis. Single-clicks on the left half of the response screen represent confidence values ranging from ‘Sure absent’ to ‘Unsure’, and the right half represent ‘Unsure’ to ‘Sure present’. As such, purple dots on the left half represent increasing confidence in the absence of a physically absent target (correct rejection), while orange dots on the left half represent confidence in the absence of a target that was physically present (miss). Orange and purple dots on the right-hand side represent, respectively, the confidence in present targets (hits), and confidence in target presence when, objectively, no target was presented (false alarm). A qualitative inspection reveals a dense diagonal cloud of responses, indicating that confidence in the presence and absence of targets correlated with attentional state ratings. This diagonal density of responses can be appreciated in Figure 2B, where the absolute value of confidence from ‘Unsure’ to ‘Sure’, is plotted against attentional state ratings, pooling both sure present and sure absent on the x-axis. To assess the strength of these correlations quantitatively, we calculated the non-parametric linear correlation coefficient between attention and confidence ratings, separately for each participant for target-present and target-absent trials. This analysis revealed a consistently positive correlation between trial-wise attentional state ratings and confidence, both in the presence of a target (one-sample *t*-tests against zero, *t*(11) =7.74, *p* < .001, *d* = 2.23) as well as the absence of a target (*t*(11) = 2.61, *p* =.025, *d* = 0.78). The strength of these correlations differed significantly, revealing an asymmetry between subjective measures of attention and decision confidence in the presence or absence of a target (paired samples t-test, *t*(11) = 5.23, *p* < .001, *d* = 1.51; Figure 2C).

In Experiment 2, participants rated the visibility of the target, responding to the prompt “How much of the target did you see?”, by clicking on the x-axis between the ranges of ‘None’ to ‘All’. As such, purple dots at the far-left value (zero) of the visibility scale represent correct rejections (trials without a target, rated as such) and orange dots at this value represent missed targets, whereas purple and orange dots with non-zero values represent false alarms and hits, respectively. In contrast to Experiment 1, no consistent correlation was observed between visibility judgements and attention ratings (*t*(8) = .09, *p* =.93; Figure 2F).

We return to this asymmetric pattern of responses in our Discussion. For now, we note two aspects of these data that provide important context for the detailed EEG analyses to follow. First, the results indicate that participants did not base their attentional state ratings solely on their sensory experience of seeing vs. not seeing a target stimulus (“I saw a target clearly so I must’ve been paying attention”, cf. (Head & Helton, 2018)): Confidence that a target was absent increased rather than decreased with attention ratings in Experiment 1, and no hint of a correlation was apparent between visibility and attention ratings in Experiment 2. Second, the contrast between Experiments 1 and 2 suggest that different information is conveyed in confidence and visibility ratings, with confidence being markedly more sensitive to variations in (rated) attentional state.

#### Matched objective performance and metacognitive sensitivity

In contrast to this varied pattern of subjective responses, objective task performance was very similar across both experiments (Figure 3). Before each experiment, target contrast was adapted using a staircase procedure to approximate 75% detection accuracy for each participant. Mean contrast values were 0.15 (*SD* = 0.02) for Experiment 1, and 0.16 (*SD* = .02) for Experiment 2. Overall accuracy, incorporating target-absent trials, was 81% (*SD* = 4%) for Experiment 1, and 80% (*SD =* 8%) for Experiment 2, with no significant difference in performance between experiments, *t* < 1 (Figure 3A). Mean detection rates in both experiments (Experiment 1 = 71%, *SD* = 8%; Experiment 2 = 73%, SD = 8 %) were not significantly different, *t* < 1. False alarm rates (Experiment 1 = 9%, *SD*= 5%; Experiment 2 = 13%, SD = 13 %) were also not significantly different, *t*(19) = −1.02, *p* = .32. We observed similar results for perceptual sensitivity (d’; Experiment 1 = 2.01, *SD* = 0.57; Experiment 2 = 1.98, *SD* = 0.72, Figure 3B) and criterion (*c*; Experiment 1 = 0.43, *SD* = 0.22; Experiment 2 = 0.36, *SD* = 0.38), which did not significantly differ between experiments (both *p* >.59).

**Figure 3.**
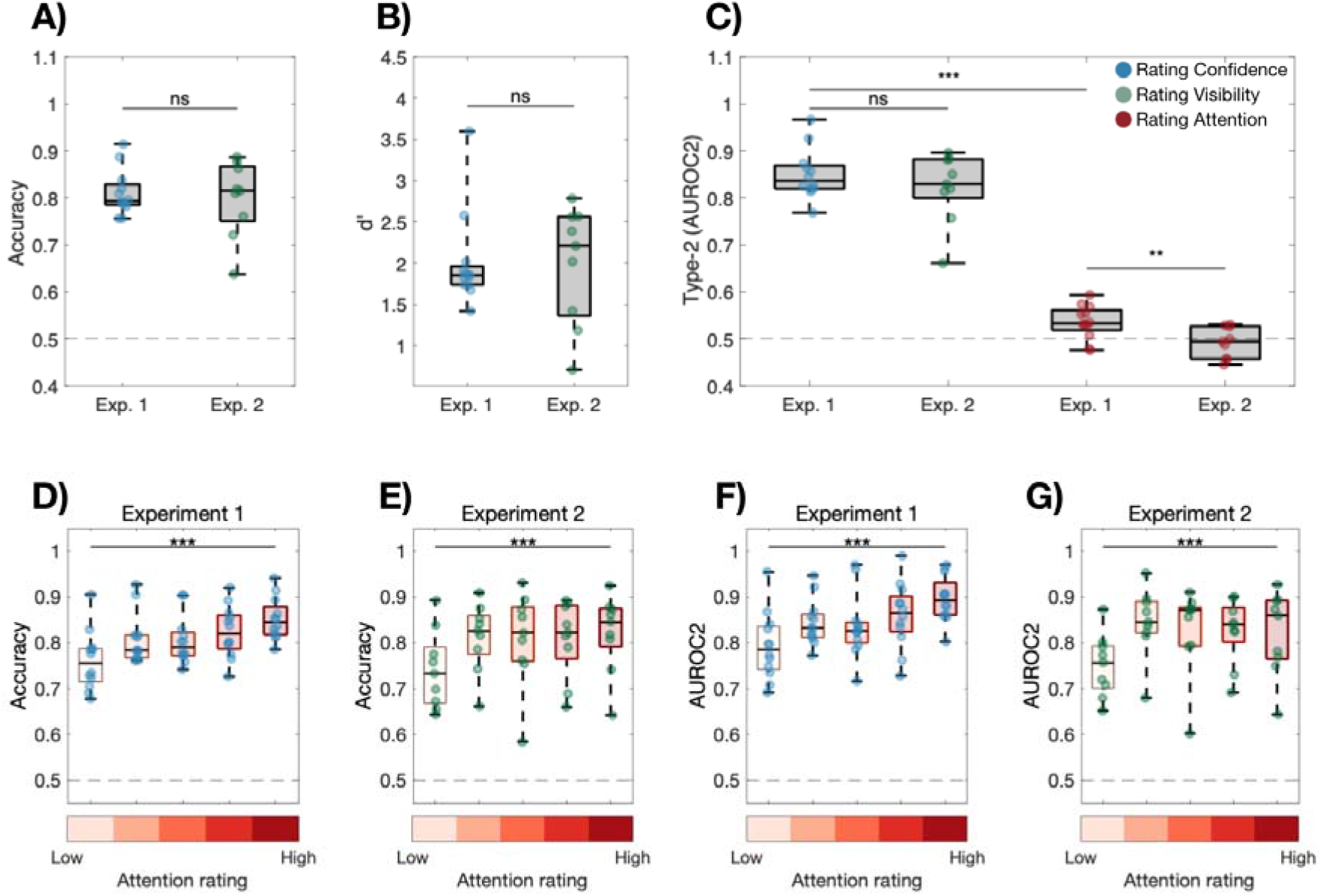
Objective and metacognitive accuracy in both experiments. A) No significant difference was observed in objective accuracy across experiments. B) Signal-detection theory measures of sensitivity (d’) were also similar across experiments. C) Metacognitive sensitivity was greatest for confidence and visibility judgements and did not differ significantly between experiments. Metacognitive sensitivity based on attention was significantly stronger in Experiment 1, although significantly weaker than metacognitive sensitivity based on confidence or visibility judgements in both experiments. D-E) In both experiments, accuracy increased with the intensity of subjective attention. F-G) In both experiments, metacognitive sensitivity also increased with subjective attention. In each box, the bottom, central, and top line indicate the 25th, 50th, and 75th percentiles respectively. Whiskers extend to the furthest data points.

We also calculated the metacognitive (type-2) sensitivity based on confidence and visibility ratings. Type-2 sensitivity captures the degree to which subjective ratings correlate with the objective likelihood of successful task performance—i.e., the degree to which the true positive rate exceeds the false positive rate at each rated value of confidence/visibility The mean type-2 performance in Experiment 1 (*M* = 0.85, *SD* = .07), was not significantly different to Experiment 2 (*M* = 0.82, *SD* = .07; *t*(19) = 1.05, *p* = .31; Figure 3C), and both differed significantly from chance (both t > 13, p <.001), despite the large differences observed in the pattern of subjective responses.

We also took the opportunity to calculate metacognitive sensitivity based on attention state ratings—i.e., the degree to which participants’ attentional state ratings were calibrated with the likelihood of a correct response. Although a nascent literature, metacognitive sensitivity based on attention ratings has recently been shown to approximate type-2 sensitivity based on confidence, in a somatosensory detection task (e.g., Whitmarsh et al., 2014; 2017). In the current visual detection tasks, type-2 sensitivity based on attentional state ratings in Experiment 1 (*M* = 0.54, *SD* = .05) was significantly lower than when based on confidence (*t*(11) = 18.74, *p* = 1.07 x 10^-9^, *d =* 5.41). Similarly, type-2 sensitivity based on attentional state ratings in Experiment 2 was significantly lower than visibility-based type-2 sensitivity (*M* = .49, *SD* = .03; *t*(8) = 11.60, *p* = 2.77 x 10^-6^, *d* = 3.87). Only the attention-based type-2 sensitivity in Experiment 1, was significantly above chance (*t*(11) = 3.51, *p* = .005, *d* = 1.01) and the strength of this type-2 sensitivity differed significantly between experiments (*t*(19) = 2.88, *p* < .01, *d* = 1.03). This latter result mirrors the patterns shown in Figure 2, in which attention ratings correlated with confidence (in Experiment 1) but not visibility ratings (in Experiment 2).

Next, we investigated whether objective accuracy and metacognitive sensitivity varied with participants’ evaluations of their attentional states (Figure 3D-G). Replicating previous findings, performance varied significantly as a function of rated attention, with objective accuracy differing significantly across attention quintiles in Experiment 1 (*F*(4,44) = 15.71, *p* < .001, *η*_p_^2^ = .59) and Experiment 2 (*F*(4,44) = 7.83, *p* < .001, η_p_^2^ = .50). Perceptual sensitivity (d-prime) increased with attention ratings in Experiment 1 (*F*(2.14, 23.54) = 6.75, *p* < .001, *η*_p_^2^ = .38; Greenhouse-Geisser corrected), but not significantly in Experiment 2 (*p* = .06). Criterion was not affected by attention in either study (*p*s > .2). A more novel finding was that metacognitive sensitivity also significantly increased alongside higher attention ratings in both Experiment 1 (*F*(4,44) = 12.09, *p* < .001, *η*_p_^2^ = .52), and Experiment 2 (*F*(4,44) = 6.67, *p* < .001, *η*_p_^2^ = .46). Thus, when more attentive, participants were not only better at the task, but also more accurately evaluated their perceptions and decisions.

Overall, therefore, in our behavioural data we observed quantitatively distinct patterns of responses when participants were asked to report either their decision confidence and attention, or visibility and attention, despite matched objective performance. In both experiments, performance increased with self-rated attention, yet only confidence, but not visibility, also positively correlated with attention ratings. To unpack this discrepancy, we turn to the strength of alpha band activity, which has been linked to the subjective intensity of visibility (Benwell et al., 2017), confidence (Samaha et al., 2017) and attention (Macdonald et al., 2011) in visual tasks.

### EEG results

We analysed the amplitude of alpha oscillations (8-12 Hz) over a 1 s preparatory period, from the onset of the words ‘Get-Ready’ to the first presentation of an image in the RSVP stream (hereafter ‘alpha amplitude’). Consistent with previous reports (Macdonald et al., 2011; Samaha et al., 2017), we observed alpha amplitude to be strongest over a cluster of parieto-occipital electrodes, and focus our remaining analysis on this subset (Figure 4). To preview our results, in both experiments, we observed that subjective attention ratings decreased with increased alpha amplitudes. In contrast to these linear effects, we observed a quadratic, inverted-U function linked preparatory alpha to subjective confidence and visibility.

**Figure 4.**
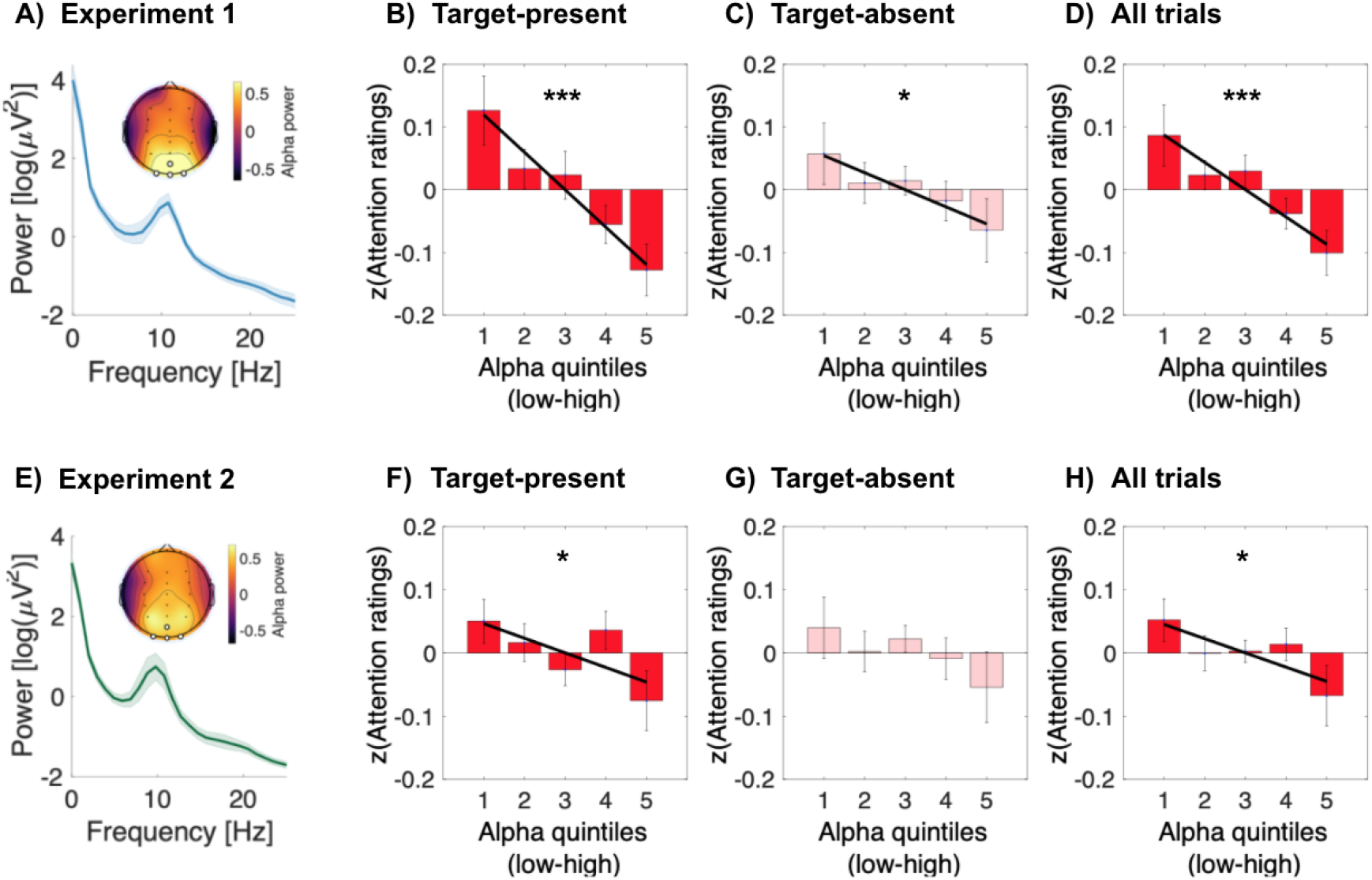
Preparatory Alpha amplitude is negatively correlated with attention ratings. In Experiment 1, strong preparatory alpha over occipital electrode sites correlated negatively and linearly with attention ratings on both target-present (B) and target-absent (C) trials, as well as in the pooled data (D). In Experiment 2, a similar topography of alpha band activity (E) negatively correlated with attention ratings on target-present trials (F) and in a pooled analysis (H), but not reliably in target-absent trials (G). Error bars represent 1 SEM, corrected for within-participant comparisons (Cousineau, 2005). Black lines display linear lines of best fit. Asterisks denote significant linear effects. *** p < .001, * p < .05.

#### Alpha amplitude is negatively correlated with subjective attention

For the effect of alpha on subjective attention ratings, when rating attention on all trials, the linear model differed significantly from the basic model, confirming a significant linear effect of alpha amplitude on subjective attention in Experiment 1 (*χ*^2^ (1) = 16.14, *p* = 5.90 x 10^-5^, β = −0.04 [−0.06, −0.02]). We further subdivided our analysis into target-present, and target-absent cases. Our motivation was to inspect whether the intervening presence (or absence) of a target within the RSVP stream would impact upon the observed relationship between preparatory alpha amplitude and attention. A key point is that this distinction allows us to investigate whether the influence of alpha exclusively biases the strength of evidence in favour of target detection.

When restricted to target-present trials, a linear effect of alpha was again the best fitting model in Experiment 1 (*χ*^2^ (1) = 19.94, *p* = 7.99 x 10^-6,^ β = −0.06 [−0.08, −0.03]). When analysing the matched subset of target-absent trials, a weaker effect of preparatory alpha on attention was observed (*χ*^2^ (1) = 5.13, *p* =.024, β = −0.03 [−0.05, −0.004]). We formally tested for the equivalence of regression coefficients (cf. Equation 4, Paternoster et al., 1998) and found the regression slopes to significantly differ between target-present and target-absent trial types (Z = −1.91, *p* = .028). This result indicates that although preparatory alpha was consistently negatively related to subjective attention, the effect of this relationship was strongest when reflecting on target-present, compared to target-absent trials.

The same pattern of results was present in Experiment 2. When considering all targets together, the linear model differed significantly from the basic model (*χ*^2^ (1) = 4.97, *p* = .025, β = −0.02 [−0.04, −0.003]). This effect was again strongest when considering target-present trials (*χ*^2^ (1) = 4.41, *p* = .036, β = −0.02 [−.04, −0.001]), as the linear model did not differ significantly from the basic model in target-absent trials (*p* = .6). However, the difference between the linear regression coefficients for target-present and target-absent classes was not significant (*p* = .42), reflecting the similar negative trend apparent in both trial types.

#### Alpha amplitude quadratically modulates confidence and visibility

In contrast to the monotonic and approximately linear relationship between alpha and attention ratings, alpha amplitude showed a quadratic relationship with the two other introspective ratings (confidence and visibility) that were recorded simultaneously with self-reported attention. In Experiment 1, a consistent quadratic trend was found, linking intermediate alpha strength to enhanced confidence that a target was present in the RSVP stream. This effect was strongest when considering decision confidence across all trials, as the quadratic model differed significantly from the basic model (*χ*^2^ (1) = 11.15, *p* = .004), and was a better fit than the linear model (*χ*^2^ (1) = 10.97, *p* = .0009, β = −0.02 [−0.03, −0.007]). The same quadratic trend was found when subdividing into the subset of only target-present trials but was not significant (*χ*^2^ (1) = 2.99, *p* = .08). On target-absent trials, alpha amplitude significantly and quadratically modulated confidence, i.e., (misplaced) confidence that a target was presented, and was a better fit than the basic (*χ*^2^ (1) =6.58, *p* = .037) and linear models (*χ*^2^ (1) =6.22, *p* = .013, β = −0.02 [−0.04, −0.004]; Figure 5A-C). In Experiment 2, when rating target-visibility, the same quadratic trend appeared. The quadratic effect was significant only on target-present trials, and a better fit than the basic (*χ*^2^ (1) =11.17, *p* = .004), and linear models (*χ*^2^ (1) =8.16, *p* = .004, β = −0.02 [−0.04, −0.007]). For target-absent trials, or when all trials were pooled together, neither linear nor quadratic models were a better fit to the data than the basic model, with only random effects per subject (all *p* > .2), reflecting very low variability in participants’ visibility ratings on target-absent trials (in which most trials were given the same [zero] visibility rating).

**Figure 5.**
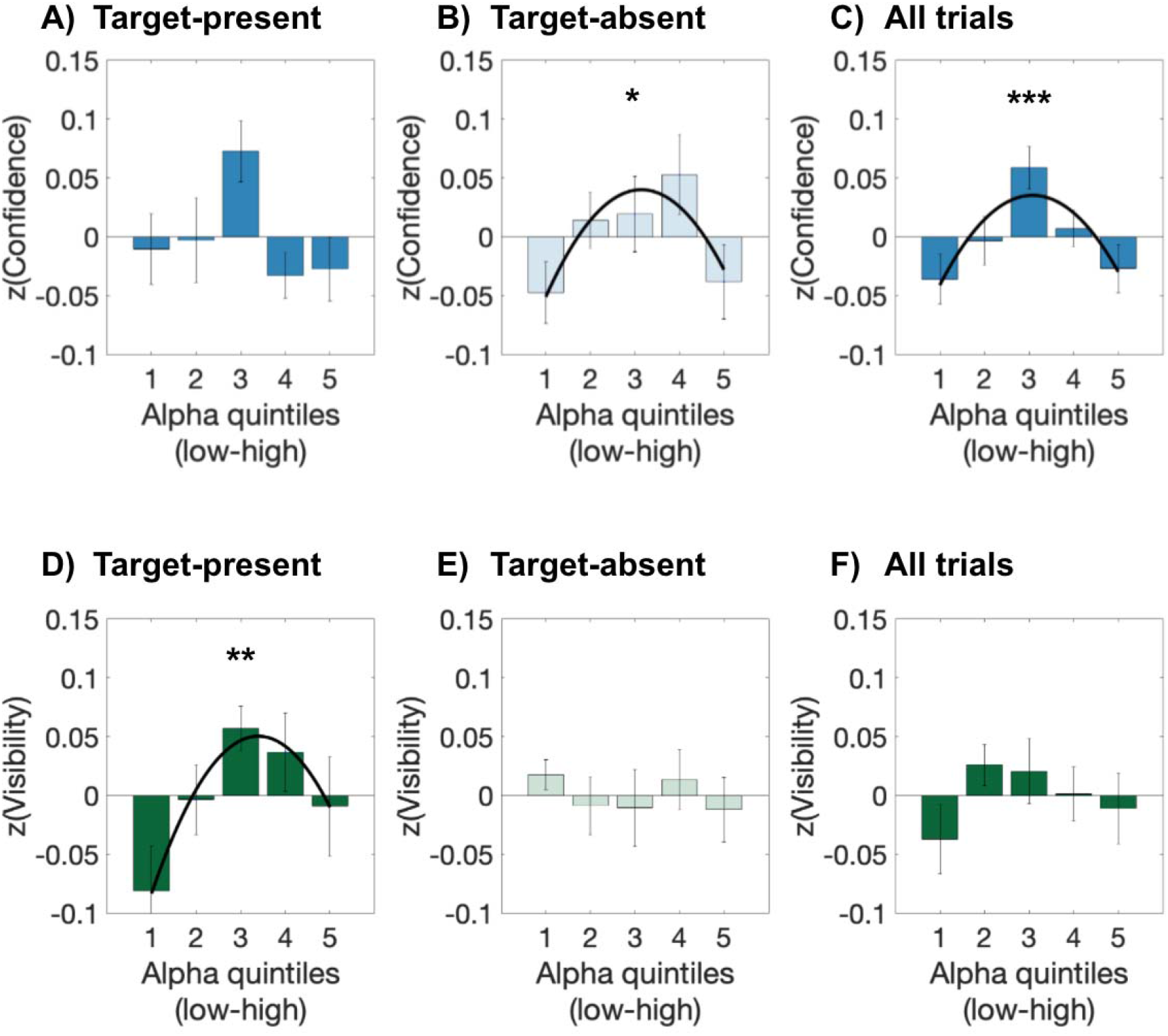
Preparatory alpha amplitude is quadratically related to subjective visibility and confidence. **A-C)** Decision confidence in the presence of a target is maximal at intermediate values of alpha amplitude. **D)** Subjective target visibility is maximal at intermediate values of alpha amplitude on target-present trials. **E)** No significant effect of preparatory alpha on visibility when targets are absent, or **F)** when pooling across all target types. Error bars represent 1 SEM, corrected for within-participant comparisons (Cousineau, 2005). Quadratic lines of best fit are shown in black. Asterisks mark significant quadratic fits. * p < .05, ** p< .01, *** p < .001

#### Alpha amplitude quadratically modulates behavioural performance

Recent work has shown that prestimulus alpha power may uniquely mediate subjective criteria, while leaving objective accuracy unchanged (for review (Samaha, Iemi, et al., 2020)). In our data, we have seen a strong and consistent relationship between the strength of alpha oscillations and subjective ratings of attentional state, as well as a significant relationship between rated attention and behavioural accuracy (Figure 3). We next examined whether alpha amplitude would also affect objective measures of performance, and focused our analyses on accuracy, hit and false alarm rates, as well as signal detection metrics of sensitivity (d’) and criterion (c). Finally, we also investigated whether metacognitive sensitivity, which was enhanced by subjective attention, would also vary with the strength of preparatory alpha amplitude. Following previous research (Busch et al., 2009; Iemi & Busch, 2018), we first normalized these responses per subject, by dividing by the mean across all alpha quintiles.

As both experiments had a very similar task structure, and objective accuracy was very similar between Experiments 1 and 2, we continued by pooling the data across all 21 participants to increase statistical power. The pattern of results we present (Figure 6) are consistent, although statistically weaker when keeping each cohort separate, as shown in the Supplementary materials.

**Figure 6.**
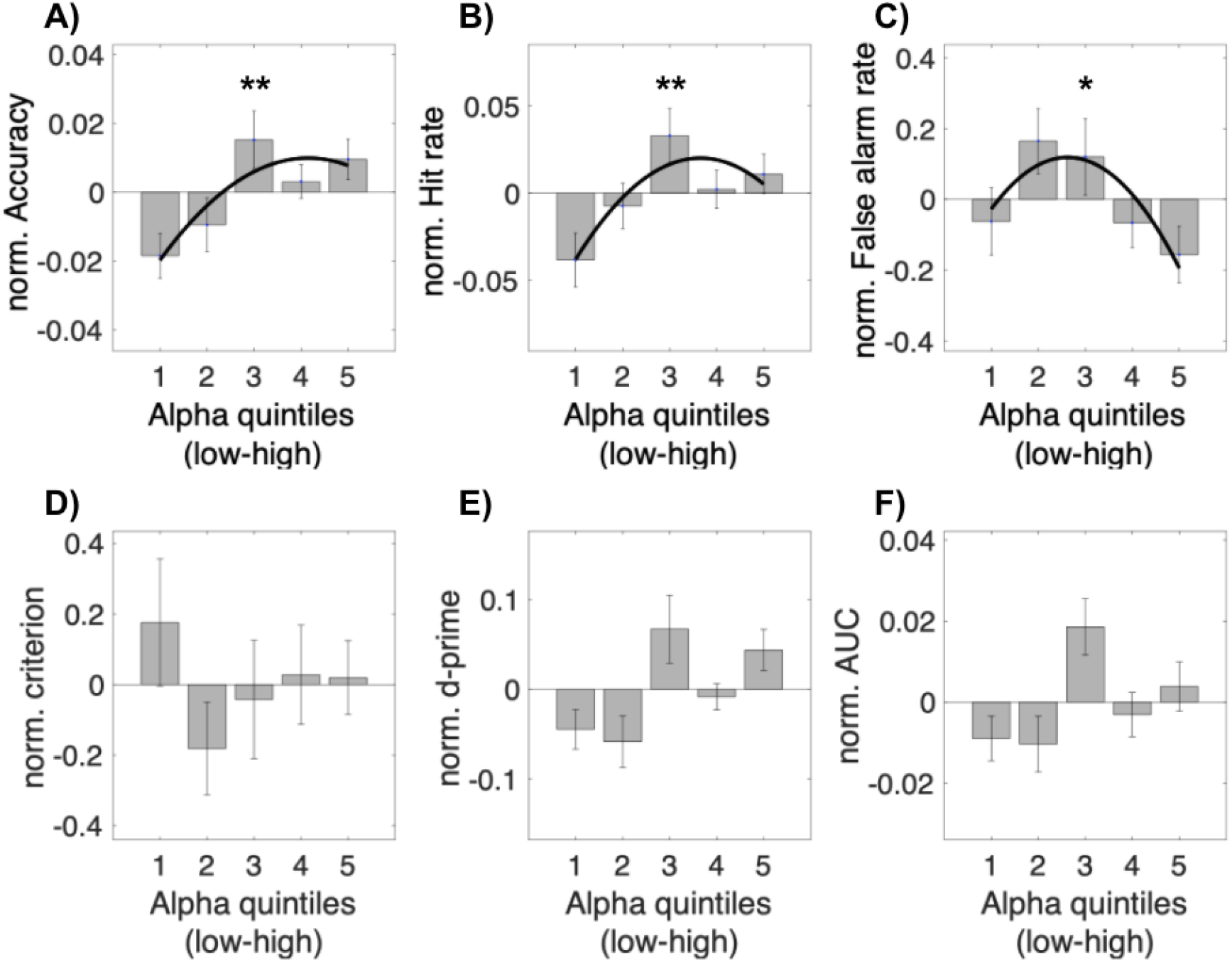
Preparatory alpha amplitude and behavioural performance. Alpha amplitude quadratically modulates a) accuracy, b) hit-rate, and c) false-alarm rate, but not d) criterion, e) d-prime, or f) AUC in combined experimental data (N=21). Responses are normalized per subject, by dividing by the mean across alpha bins, and zero centred by subtracting by 1. * p < .05 ** p < .01. For separate experiments, see Supplementary Figure 3.

Alpha amplitude in the preparatory window significantly affected overall accuracy. Both the linear (χ^2^(1) =8.40, *p* = .004, β = 0.006 [0.002, 0.01]) and quadratic models (χ^2^(1) =10.87, *p* = .005, β = −0.002, [−0.006, −0.0007]) were superior fits than the basic model. When comparing the linear and quadratic fits, neither were a better fit to the data (*p* = .12). Post-hoc comparisons, adjusting for a family-wise error rate of 10, revealed that only the lowest and intermediate alpha bins differed significantly (Bin 1 vs. 3: *t*(20) = −2.91, *p*_bonf_ = .047, *d* = −0.57). Therefore, like subjective visibility and confidence ratings, the effect was an enhancement of objective accuracy at intermediate alpha amplitudes.

In stimulus detection tasks, accuracy measures can be influenced by both the likelihood of detecting a present target, as well as withholding responses on target-absent trials. To parse these effects, we also analysed SDT stimulus-response categories of performance. Alpha amplitude significantly affected the normalized hit-rate during all trials, and a quadratic model was again the best fit to the data (χ^2^(1) =12.39, *p* = .002, β = −0.008 [−0.015, −0.001]). When comparing linear and quadratic models, likelihood ratio tests revealed the quadratic model was a significantly better fit (χ^2^(1) =5.54 *p* = .02), with post hoc comparisons again revealing that this effect was driven by a significant difference between the lowest and intermediate alpha bins (Bin 1 vs. 3: *t*(20) = −3.39, *p*_bonf_ = .011, *d* = −0.61). A quadratic model was the best fit to the data for the FA rate (χ^2^(1) =7.43, *p* = .024, β = −0.06 [−0.1, −0.008]), which significantly improved upon the linear model (χ^2^(1) =6.37, *p* = .012). Given this parallel increase in hits and false alarms at intermediate levels of preparatory alpha, it is not surprising that we do not find a significant effect of preparatory alpha amplitude on sensitivity (d-prime), somewhat in contrast to the quadratic effects apparent in the simpler measure of overall accuracy (which in our data is primarily driven by hit rate because of the low incidence of false alarms). More surprisingly, given the increase we observed in both hits and false alarms at intermediate levels of alpha, and given recent evidence that low prestimulus alpha power is associated with a more liberal detection criterion (Samaha, Iemi, et al., 2020), we found no significant effect of preparatory alpha amplitude on criterion (*p*s > .5). Similarly, alpha amplitude did not significantly affect type-2 sensitivity (*p*s > .09; Figure 6).

#### Alpha amplitude quadratically modulates event-related potentials

Across the two experiments we have observed an interaction between preparatory alpha amplitude and subjective ratings of attention, confidence, and visibility. Moreover, a dependence on the trial-type, whether targets were physically present or absent from the intervening trial-window, also modulates these effects. For example, in Experiment 1, the relationship between alpha and attention was significantly greater on target-present trials. Similarly in Experiment 2, intermediate alpha amplitudes quadratically modulated subjective target visibility, yet only when targets were physically present. Given these interactions, we hypothesised that alpha would affect the underlying neural response to target stimuli, particularly at intermediate levels of alpha amplitude . We directly tested for this relationship by focusing on two ERP measures, the P1 which reflected the initial sensory response to the RSVP stream onset, and the CPP to target stimuli embedded in half of the RSVP streams. Again, to increase statistical power, and given the identical structure of the tasks in terms of stimulus presentation, we again pooled the data across all participants for these analyses.

#### Quadratic modulation of early sensory-evoked response (P1)

How the generation of sensory evoked potentials are influenced by prestimulus neural activity is the focus of ongoing research (Gruber et al., 2014; Iemi et al., 2019; Min et al., 2007). Notably, a quadratic, inverted-U function such as the type we report above, linking preparatory alpha amplitude to confidence and visibility reports, has also been reported to link prestimulus alpha power and the amplitude of the early P1 component of the ERP (Rajagovindan & Ding, 2011). Accordingly, we tested whether the amplitude of the P1 component evoked 80-160 ms after RSVP onset was also modulated by preparatory alpha. The quadratic model was a significant improvement upon the basic (χ^2^(1) =9.47, *p* = .009, β = −0.08 [−0.15, −0.02]), and the linear model (χ^2^(1) =7.26, *p* = .007) demonstrating that alpha amplitude quadratically modulates the amplitude of the early P1 component. The same pattern, although statistically weaker, was observed in the data for Experiment 1 when analysed separately (quadratic: χ^2^ (1) =12.49, *p* = .002, β = −0.12 [−0.20, −0.03]; comparison: χ^2^ (1) =^2^ 7.36, *p =* .006), though did not reach significance in Experiment 2 (*p*s > .3) (Figure 7B).

**Figure 7.**
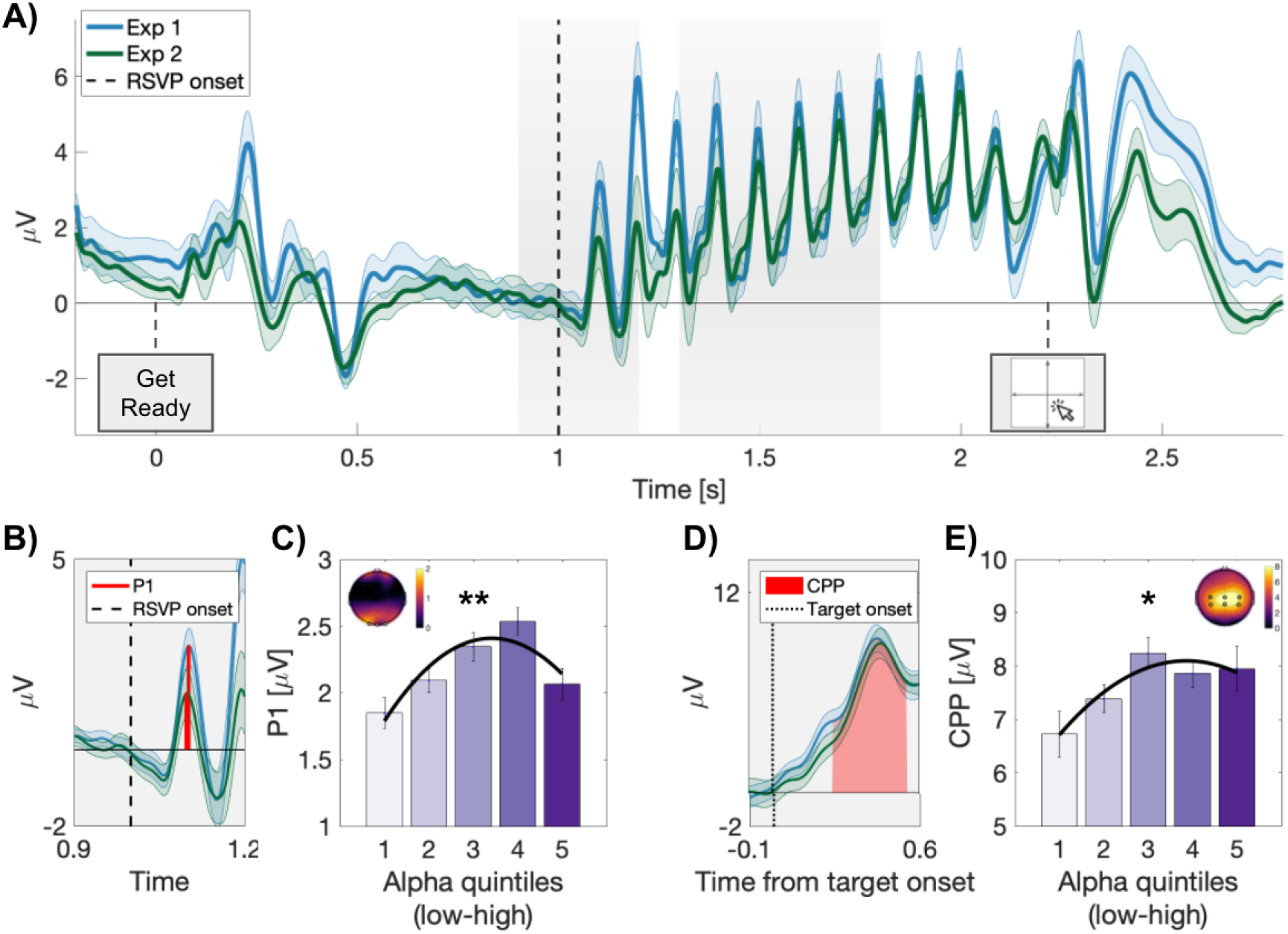
Preparatory alpha amplitude quadratically modulates event-related potentials. A) Grand average whole-trial epochs for Experiments 1 and 2. Grey shaded regions note the time windows used to calculate the P1, and target-locked centro-parietal positivity (CPP; see Methods). B) Grand average P1 from Experiments 1 and 2. C) Preparatory alpha quadratically modulates the amplitude of the early P1 component, evoked by the first image in our RSVP stream in Experiment 1. D) Grand average target locked CPP. Red shading indicates 250-550ms relative to target onset. E) Average CPP amplitude over the period 250-550 ms relative to target onset in Experiment 1. In all plots error bars and shading indicate 1 SEM, corrected for within-participant comparisons (Cousineau, 2005). * p < .05, ** p < .01.

#### Quadratic modulation of the centro-parietal positivity (CPP)

Next, as an index of decision-related processes, we investigated whether the amplitude of target-locked activity evoked on ‘Hit’ trials (successful detection of present targets) was also modulated by preparatory alpha. In the scalp EEG, we observed a typical broad CPP after target onset that was strongest over central electrodes (C3, Cz, C4, CP3, CPz, CP4). We computed the average CPP amplitude across these electrodes, over the period 250 to 550 ms relative to target onset, based on quintiles of preparatory alpha amplitude. We observed that a quadratic fit was the best fit to the data, and a significant improvement upon the basic (χ^2^(1) =6.78, *p* = .034, (β = −0.15 [−0.33 −0.03]), but not the linear model (*p* = .1). When examining each experiment in isolation, the same pattern was only significant in Experiment 1 (quadratic; χ (1) = 7.64, *p* = .02 β = −0.24, [−0.52, −0.03]), with neither the linear nor quadratic models reaching significance in Experiment 2 (*p*s > .7).

### The CPP positively correlates with subjective confidence and visibility

We have shown that alpha amplitude quadratically modulated subjective confidence and visibility, as well as the strength of early (P1) and late (CPP) event-related potentials. Previous research has also shown that the amplitude of the CPP captures the strength of a perceptual experience (Tagliabue et al., 2019), consistent with the notion that it indexes the strength of accumulated evidence in favour of a particular perceptual decision (Murphy et al., 2015; O’Connell et al., 2012; Twomey et al., 2015). We therefore next tested whether CPP amplitude in our paradigm varied with subjective ratings of confidence, visibility, or attention. Consistent with our expectations, we observed that the amplitude of the CPP varied strongly and consistently with both confidence and visibility ratings. In Experiment 1, CPP strength increased with subjective confidence (linear: (χ^2^(1) =14.13, *p* < .001, β = 0.85, [0.55, 1.15]). In Experiment 2, CPP strength also increased with subjective visibility (linear: (χ^2^(1) =14.13, *p* < .001, β = 0.85, [0.55, 1.15]).

In contrast to the consistent monotonic, linear relationship between CPP amplitude and confidence/visibility ratings, a more complex relationship was observed between CPP amplitude and attention ratings (Figure 8). In Experiment 1, although we observed that CPP amplitude was maximal at highest ratings of attention, the best fit to the data was a quadratic model rather than a linear one (quadratic: (χ^2^(1) =15.16, *p* < .001, β = 0.85, [0.55, 1.15]). By comparison, in Experiment 2, attention did not significantly predict CPP amplitude (*p*s> .43). A straightforward implication of these findings is that they provide further evidence that participants’ attention ratings do not simply reflect the strength of their perceptual experience. The specific, detailed pattern is more complex to explain. The quadratic relationship apparent in Experiment 1 would be predicted if CPP amplitude reflected the strength of evidence needed for a participant to decide that a target was present in a particular trial, given higher baseline evidence at intermediate levels of alpha (as suggested by the ERP results) and a fixed response criterion (as suggested by our SDT analysis). However, we would expect a similar relationship to hold in Experiment 2. The contrast across experiments suggests that the nature of the decision made by participants influenced the CPP, which would be consistent with this component indexing a high-level, decision-related process.

**Figure 8.**
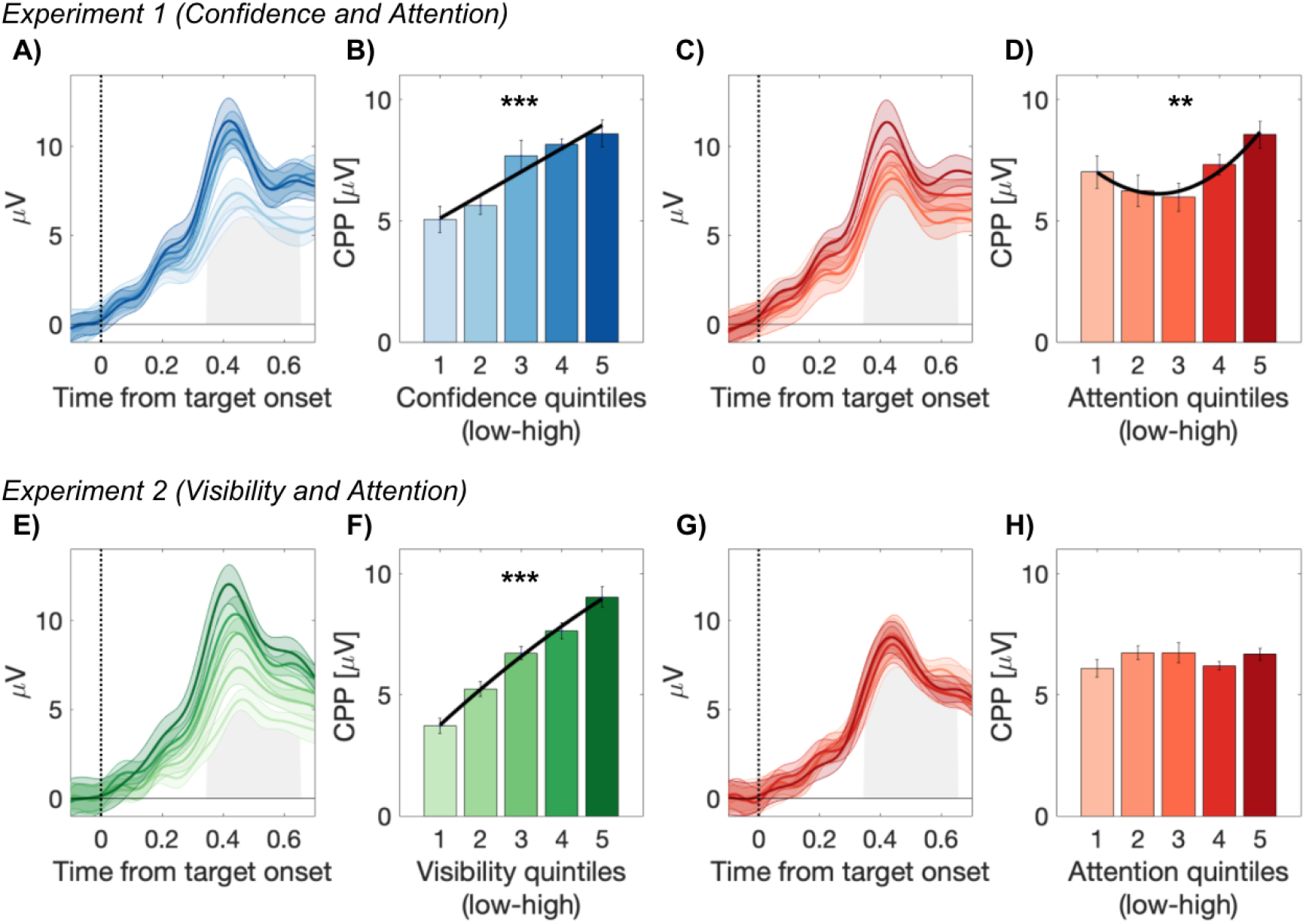
The subjective correlates of the centro-parietal positivity. CPP amplitude increases with reported confidence (A-B), and visibility (E-F), in Experiments 1 and 2, respectively. CPP amplitude also varied as a function of subjectively rated attention in Experiment 1 (C-D), but not in Experiment 2 (G-H). Grey shaded regions note 250-550 ms relative to target-onset, used to calculate the CPP.

## Discussion

This study aimed to characterise the relationship between preparatory alpha amplitude and subjective ratings of attention, confidence and stimulus visibility, and thereby provide insight into the basis of these introspective judgments. Previous work, focusing mainly on visual discrimination tasks, has demonstrated a negative linear relationship linking prestimulus alpha power to all three of these subjective criteria. Here we demonstrate that in a visual detection task, preparatory alpha amplitude does negatively correlate with subjectively rated attention, but that it quadratically modulates decision confidence and visibility. In support of this quadratic relationship, we also found that alpha amplitude quadratically modulates objective performance, as well as the amplitude of event-related potentials elicited by task stimuli. Importantly, we outline the neural commonalities and dissociations of these overlapping subjective criteria.

### Preparatory alpha amplitude and subjective reports

Given the natural correlation between cortical excitability, attention, and subjective judgements of visibility and confidence, in many situations their inter-relationships are difficult to disentangle. The present dataset is interesting in this regard because we observe that alpha amplitude showed a different relationship with attention ratings vs. ratings of confidence and visibility. Specifically, after splitting alpha amplitude into quintiles, we observed the expected negative and monotonic relationship between alpha and subjectively rated attention, but found that intermediate levels of alpha corresponded to the highest subjective ratings of decision confidence and visibility. Intermediate alpha amplitude was also associated with increased accuracy, as well as increased amplitude of early (P1), and late (CPP) sensory evoked potentials.

This inverted-U function is in contrast to recent examples of a negative and linear relationship between prestimulus alpha power and various performance measures in discrimination tasks (Benwell et al., 2017; Iemi et al., 2017; Samaha et al., 2017). Our aim here was not to explore the mechanisms underpinning this quadratic relationship: Rather, observing this effect gave us the opportunity to dissociate measures – of attentional state, evoked responses, task performance, and performance evaluations – that are typically mutually correlated. However, it is interesting to ask what may drive such a quadratic association. A quadratic link between prestimulus oscillatory power and performance has previously been reported in somatosensory detection tasks (Linkenkaer-Hansen et al., 2004; Zhang & Ding, 2010), and between alpha power and the amplitude of early visually evoked potentials (Rajagovindan & Ding, 2011). In their model, Rajagovindan and Ding (2011) proposed that the total output of a neural ensemble can be characterised by its position on a sigmoidal curve, with each point on the curve being jointly determined by background synaptic activity and the addition of a sensory evoked response (see Rajagovindan & Ding, 2011, for details). Their model predicts maximal sensory-evoked output at intermediate levels of alpha power, where the sigmoidal curve is steepest, and was supported by measuring the amplitude of the P1 response at attended, compared to unattended locations. Our visual detection tasks differ in many important ways, yet we also find that early visual evoked responses in the P1 window are quadratically modulated by the strength of alpha oscillations. As an extension of these results, here we can add that subjective visibility and confidence are also greatest at intermediate levels of preparatory alpha amplitude. Interestingly, in all cases, increased confidence/visibility that a target was present was associated with intermediate alpha, even on exclusively target-absent trials (Figure 5A-C). Thus, the relationship between preparatory alpha amplitude and decision confidence appears to be directional: Intermediate alpha amplitude does not enhance confidence in any decision, but enhances confidence in perceiving the presence of a target, even on exclusively target-absent trials.

The present work complements recent evidence linking the amplitude of prestimulus oscillations to the intensity of subjective reports (Samaha, Iemi, et al., 2020), by clarifying the role of intervening event-related potentials on measures of confidence, visibility and attention. Previous links between prestimulus power and subjective reports studied each in isolation, or omitted ERP analyses (Benwell et al., 2017; Samaha et al., 2017; Whitmarsh et al., 2021), which in the present work have revealed novel dissociations between these overlapping subjective criteria. Specifically, we find that alpha oscillations during task preparation negatively correlate with the intensity of subjective attention on both target-present and target-absent trials, and when rating either visibility or confidence in the intervening trial window. Participants could distinguish these fluctuations in attention from the strength of sensory evidence when rating perceived target visibility, which positively correlated with the amplitude of sensory evoked responses, whereas ratings of attention did not (CPP cf. Figure 8). In contrast, confidence ratings incorporated both the context of attentional state and the strength of sensory evidence, as these subjective reports were positively correlated, and increased concomitantly with CPP amplitude.

We also observed a quadratic relationship linking the strength of alpha oscillations to both the amplitude of event-related potentials and the strength of visibility and confidence judgements. We can now characterise the information that underpins subjective reports of attention and confidence in this way: preparatory alpha amplitude is negatively correlated with the intensity of subjective attention, and quadratically modulates the strength of sensory-evoked potentials. The strength of these sensory-evoked potentials, in turn, partially determine the intensity of subjective visibility and confidence-with the latter also incorporating, and correlating, with the intensity of subjective attention. These observations add to a growing literature that the CPP represents the accumulation of decision likelihood based on internal states, which include the subjective certainty of a decision (Gherman & Philiastides, 2015; Rangelov & Mattingley, 2020; Tagliabue et al., 2019), as opposed to a pure index of physical sensory evidence (O’Connell et al., 2012). One caveat however, is that interpreting the relationship between alpha and evoked-responses is complicated by recent work showing that alpha oscillations have a non-zero mean (Iemi et al., 2019). This asymmetry in mean voltage could result in the quadratic effect reported here being a mixture of this asymmetry and a true modulation of the evoked responses.

The inverted-U function is in contrast to recent examples of a negative and linear relationship between prestimulus alpha power and detection performance (e.g. (Iemi et al., 2017) as well as confidence and visibility in 2AFC visual discrimination tasks (e.g. (Benwell et al., 2017; Samaha et al., 2017). As such it is important to consider important differences between our present and previous works which may contribute to these discrepancies. Most notably, discrimination and detection judgements may be supported by fundamentally distinct processes, and recent work has begun to describe independent behavioural (Kanai et al., 2010; Meuwese et al., 2014), as well as neural correlates (e.g. Mazor & Fleming, 2020) that distinguish these judgement types. More practically, detection in the present work required identification of a single image within an RSVP stream, and alpha oscillations were measured in a preparatory window after the instruction to “Get Ready” was displayed on screen. As a result, decision making involved processing target signals embedded in noise, and thus integrating evidence over an extended period. This is in stark contrast to previous examples that usually employ short duration, near threshold targets on an otherwise unchanged background (e.g., Iemi et al., 2017). RSVP streams also began after a fixed interstimulus interval, and targets were predictably located at 50 ms intervals within this stream. These features change the anticipatory and predictive demands of our paradigm compared to previous work, and it is presently unclear how these differences may combine to interact with alpha power and target detection (Clayton et al., 2015, 2018; Van Diepen et al., 2019).

In addition to the proposed link between alpha amplitude and perceptual performance, another non-exclusive possibility is that the phase of alpha oscillations rhythmically modulate inhibition-excitation cycles, which also determine perceptual outcomes (Chapeton et al., 2019; Jensen et al., 2012; Klimesch, 2012; Klimesch et al., 2007; Mathewson et al., 2012; Mazaheri & Jensen, 2010). For example, it has previously been reported that the phase of prestimulus alpha oscillations can determine whether near-threshold targets are detected (Busch et al., 2009; Mathewson et al., 2009). Moreover, the phase of spontaneous alpha can be adjusted under top-down control, in anticipation of stimulus onset (Samaha et al., 2015); yet see (van Diepen et al., 2015)). To our knowledge, however, whether subjective estimates, such as confidence, visibility, or attention are also modulated by anticipatory phase have not been reported. Although it is beyond the scope of the present manuscript, it was clear in our dataset that subjective confidence and the visibility of a target were systematically biased by the phase of alpha during task preparation, while attention was not (Supplementary Figure 4). Future work will be necessary to untangle these complex relationships, and further determine how the phase of alpha may similarly mediate sensory-evoked potentials.

### The relationship between attention, confidence, and visibility ratings

Our findings provide new insight into the relationship between introspective reports of attention and sensory experience. Although attention and confidence have traditionally been studied in isolation, recent research has begun to expand our understanding of their relationship. Predominantly, this has been achieved by contrasting confidence between attended and unattended conditions. For example, when spatial attention is validly cued toward a target location, subjective confidence increases in discrimination tasks compared to confidence at unattended, or invalidly cued locations (Kurtz et al., 2017; Zizlsperger et al., 2012, 2014); yet see (Wilimzig et al., 2008), for the opposite effect). As a complement to these effects of cued attention, here we show that increased subjective attentional engagement in a task is also associated with increases in confidence in a graded manner. The intensity of attention also increased both objective performance accuracy, and metacognitive sensitivity in our paradigm. As a consequence, our results speak to the value of monitoring subjective attentional demand in perceptual research because even matched conditions, if differing in perceived attentional effort, will result in significant differences to both subjective and objective performance.

The incorporation of attention-related information may improve perceptual decisions by reducing uncertainty (Denison et al., 2018), or alternatively, by boosting confidence due to an apparent increase in stimulus contrast (Carrasco et al., 2004). Indeed, perceptual confidence has been tightly yoked to the amount of sensory information that is available in favour of a decision (for review; (Mamassian, 2016)). In this regard, the effects of attention are reminiscent of the near-ubiquitous effect of objective task difficulty on confidence, whereby easier tasks are associated with greater confidence in correct responses and reduced confidence in errors, and therefore an overall increase in metacognitive sensitivity (Kepecs & Mainen, 2012; Maniscalco & Lau, 2012). However, our results suggest that attention does not only affect confidence indirectly via changes in signal quality. If so, we might not expect significant effects of attention on decision confidence in the absence of a target (which were clearly apparent in Experiment 1), and we would expect similarly strong effects of attention on visibility judgments (which were not observed in Experiment 2).

Instead, confidence reports appear to integrate information about attentional state more directly such that, above and beyond any effects of attention on signal quality, people experience or express higher confidence in decisions they make when focused on (vs. distracted from) the task at hand. Thus, confidence correlates strongly with attention, more so than visibility ratings, and in a manner that can be normatively justified: Intuitively, one should place less trust in a given perceptual impression (whether of presence or absence of a target) when it is derived from an inattentive glimpse than from careful focused viewing. This interpretation is consistent with other recent suggestions that confidence is not a direct readout of accumulated evidence strength, but instead integrates relevant contextual information (Bang et al., 2017; Boldt et al., 2017; Kiani et al., 2014). Such a two-stage model of confidence formation (cf. (Shekhar & Rahnev, 2018) is in contrast to earlier proposals that confidence directly reflects the strength of accumulated evidence (for reviews; (Pleskac & Busemeyer, 2010; Yeung & Summerfield, 2012) but aligns with other evidence that confidence can be manipulated without a change in sensory evidence (e.g. (Cortese et al., 2016, 2017). This higher-order influence on decisions (see Denison et al., 2018; Mazor et al., 2020; for related discussions), may have been exacerbated in our task paradigm, as responses were not speeded, allowing sufficient time for reflection and adjustment of subjective ratings between the RSVP stream and response options. Future work will be necessary to test whether reduced stimulus-response intervals mediate the correlation between target-absent confidence and attention ratings.

This correlation between attention and confidence notwithstanding, the two ratings showed clear dissociations: Confidence showed a linear relationship with the strength of sensory evidence as reflected in sensory evoked potentials but varied quadratically as a function of preparatory alpha amplitude, whereas attention ratings showed the opposite pattern. More broadly, we found little evidence that attention ratings are inferred indirectly from the strength of perceptual evidence accumulated for a decision (“I saw a target clearly so I must’ve been paying attention”, cf. (Head & Helton, 2018), and instead they seem to depend on more direct insight into the true underlying attentional state (as it is reflected in alpha amplitude, for example). This insight might come from monitoring the state of sensory systems themselves, but perhaps more plausibly derives from access to one’s current level of motivation and effort expended on the task (i.e., information about the strength of exerted attention and control). That said, a nuance of the present results was that participants’ attention ratings differed subtly across experiments, for example showing a stronger relationship with CPP amplitude in Experiment 1 than Experiment 2. While it remains possible that comparisons between these groups are hampered by differences in statistical power, another possibility is that the specific wording used for the visibility question in Experiment 2 (‘How much of the target did you see?’) may have primed a quantitative, as opposed to qualitative use of the visibility scale, and encouraged participants to distinguish their sensory experience from subjective level of engagement in the task. In contrast, the experiential focus of the confidence question in Experiment 1 (“How confident are you?”) may have led participants to base their attention ratings more on experiential cues such as the strength of their perceptions (e.g., it would be counterintuitive to indicate you were sure a target was present/absent even though you had been paying little attention to the task). Although speculative, this possibility can easily be tested in future research, by adapting the visibility prompt to instead include a qualitative estimate of perceptual awareness that is a standard in consciousness research (e.g. “How clear was your visual experience?”; see (Overgaard & Sandberg, 2012; Ramsøy & Overgaard, 2004; Sandberg et al., 2010)).

## Conclusion

Our study sheds new light on the interaction between preparatory alpha amplitude and subjective phenomena in an RSVP target detection task. Alpha amplitude negatively and linearly correlated with the intensity of subjective attention, yet quadratically modulated the strength of decision confidence and visibility. This partial independence speaks to the importance of choosing appropriate subjective response options in experimental tasks, and for future studies of metacognition, suggesting that confidence reports (but not visibility) may conflate attentional state ratings. Importantly, understanding the influence of alpha on subjective criteria can be enriched by considering the intervening effect of alpha on stimulus-evoked responses. We show that people are able to distinguish and separately report their sensory experience (here: stimulus visibility) and their attentional state, with the former reflected in sensory-evoked potentials and the latter in alpha oscillations. But they appear to combine these signals when they report the reliability of their perceptions as reflected in the confidence they express in their decisions. Collectively, these findings provide insight into the commonalities and dissociations among different subjective reports in their psychological properties and neural underpinnings.

## Acknowledgements

The authors acknowledge support of the UK Ministry of Defence through the Defence Science and Technology Laboratory (DSTLX 1000128890), under the Bilateral Academic Research Initiative programme.

## Data and Code Availability

All raw data and analysis code have been uploaded to a repository on the Open Science Framework (https://osf.io/j2cah/).

## Supplementary Methods

For our phase-based analysis, our analysis focused on whether subjective criteria varied as a function of alpha phase angle. For this analysis, we calculated complex-values using our FFT transform, as described above, and now retained the single-trial phase angles. For each participant, we next split the single trial phase angles into 11 phase bins and averaged the subjective criteria of interest within each bin. As the objective preferred phase angle can vary across individuals, we first centred on each individual’s preferred phase angle, based on the maximum subjective criteria, before averaging across subjects. As in previous uses of this analysis (e.g., Busch et al., 2010), a peak at the preferred phase angle is trivial, because of this realignment. A significant effect of phase on subjective criteria can only be inferred after omitting this central phase bin and was tested for using repeated-measures ANOVA.

## Supplementary Figures

**Supplementary Figure 1.**
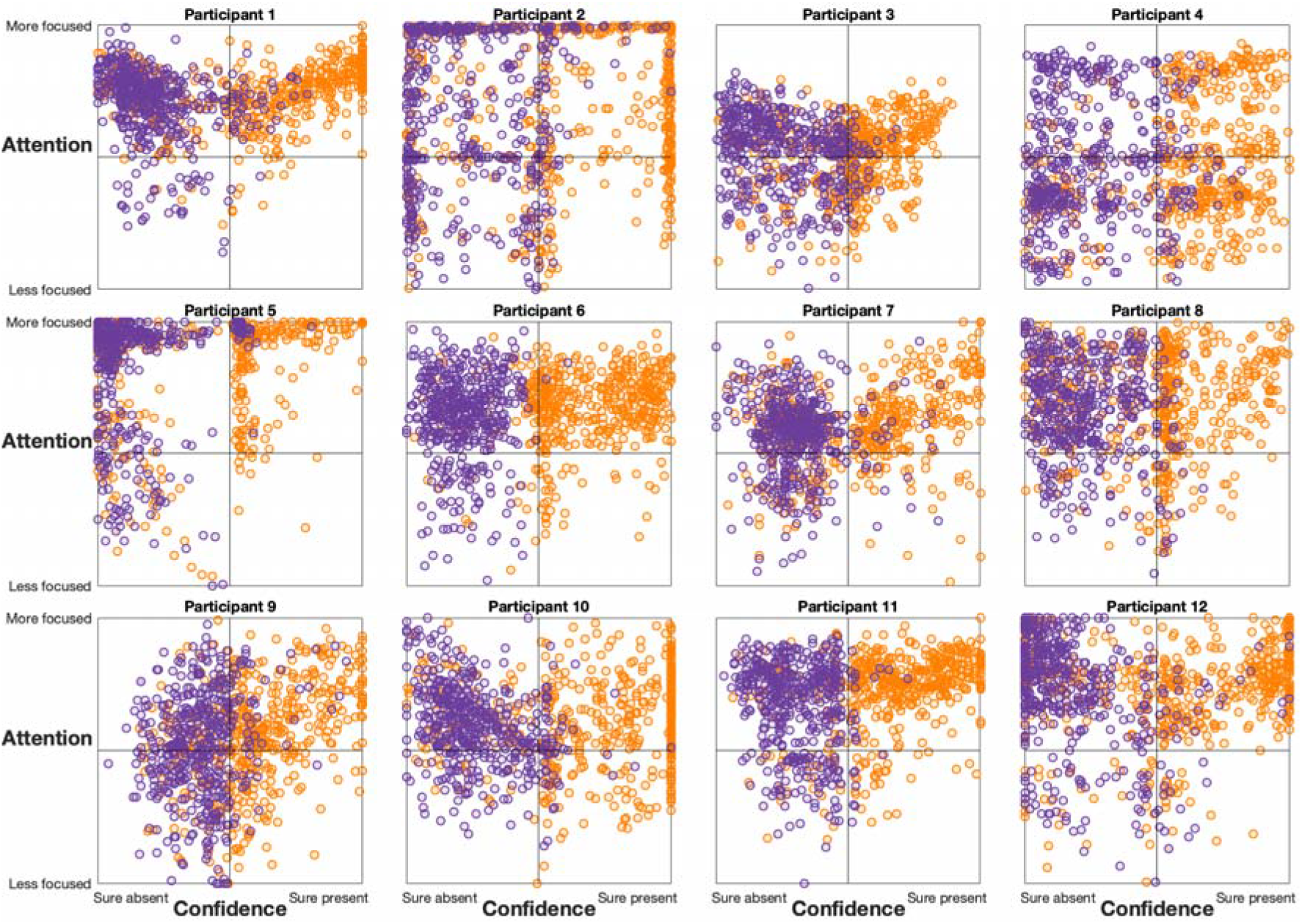
Behavioural responses for participants in Experiment 1.

**Supplementary Figure 2.**
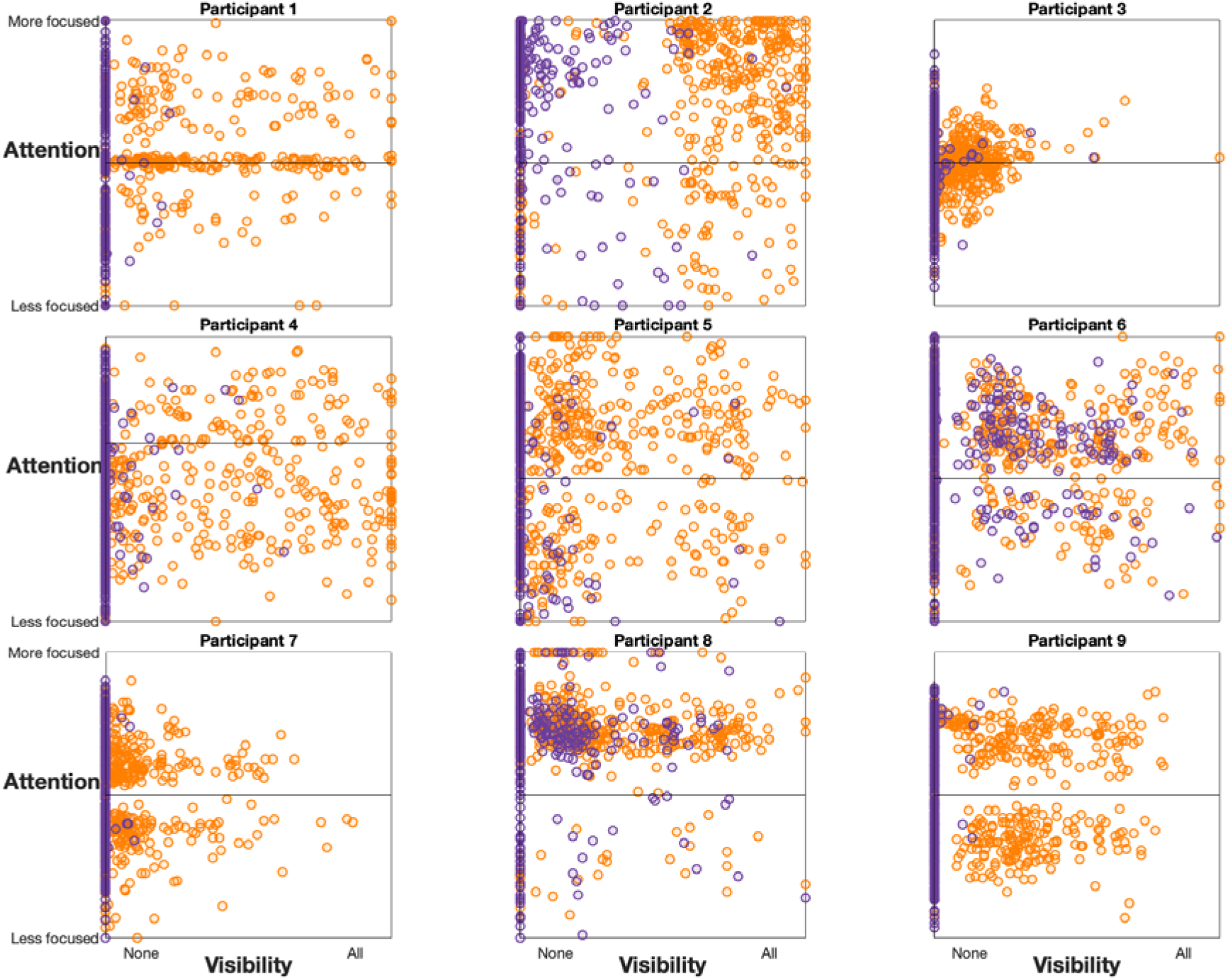
Behavioural responses for participants in Experiment 2.

**Supplementary Figure 3.**
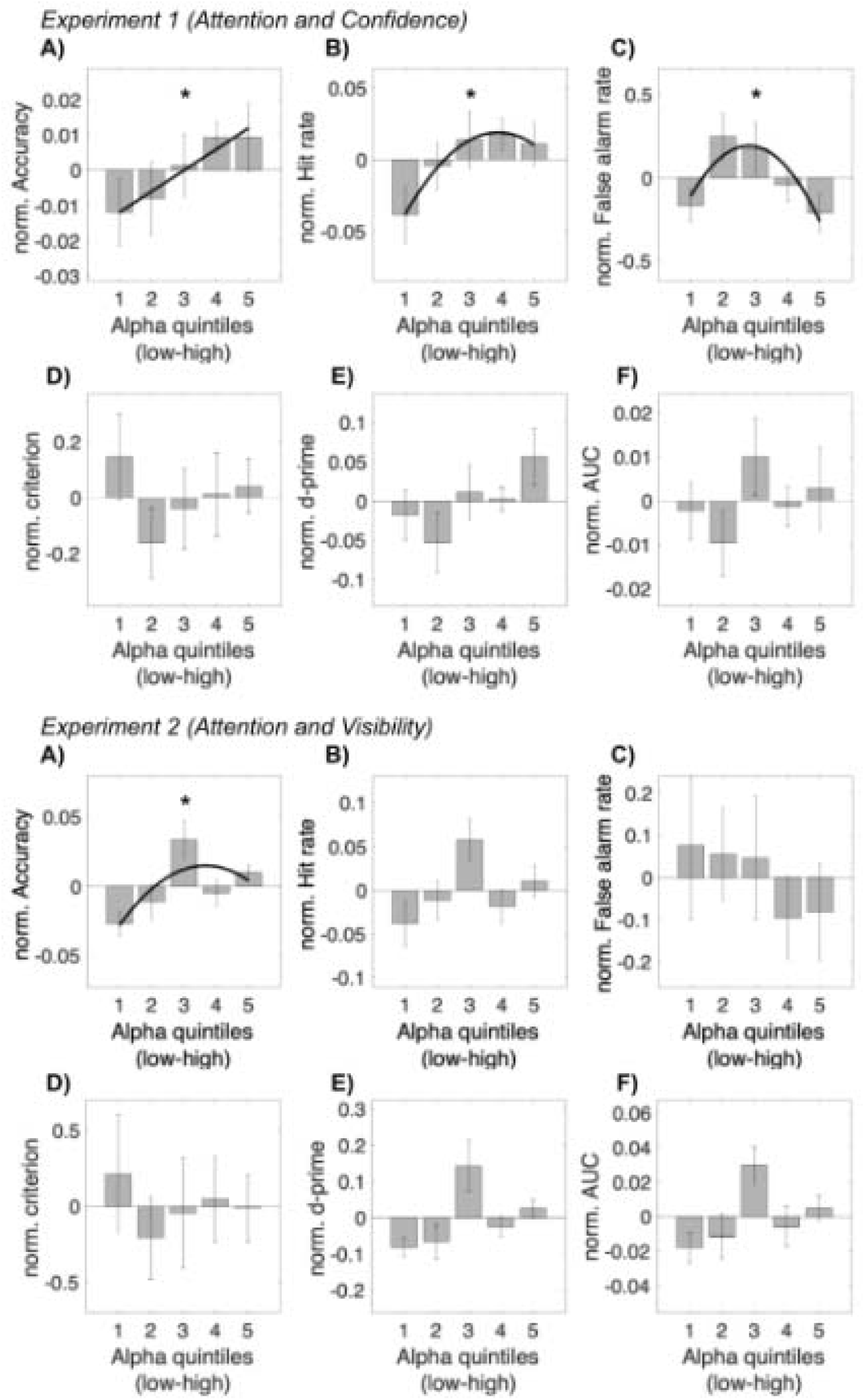
Alpha amplitude and objective performance in separate experiments.

**Supplementary Figure 4.**
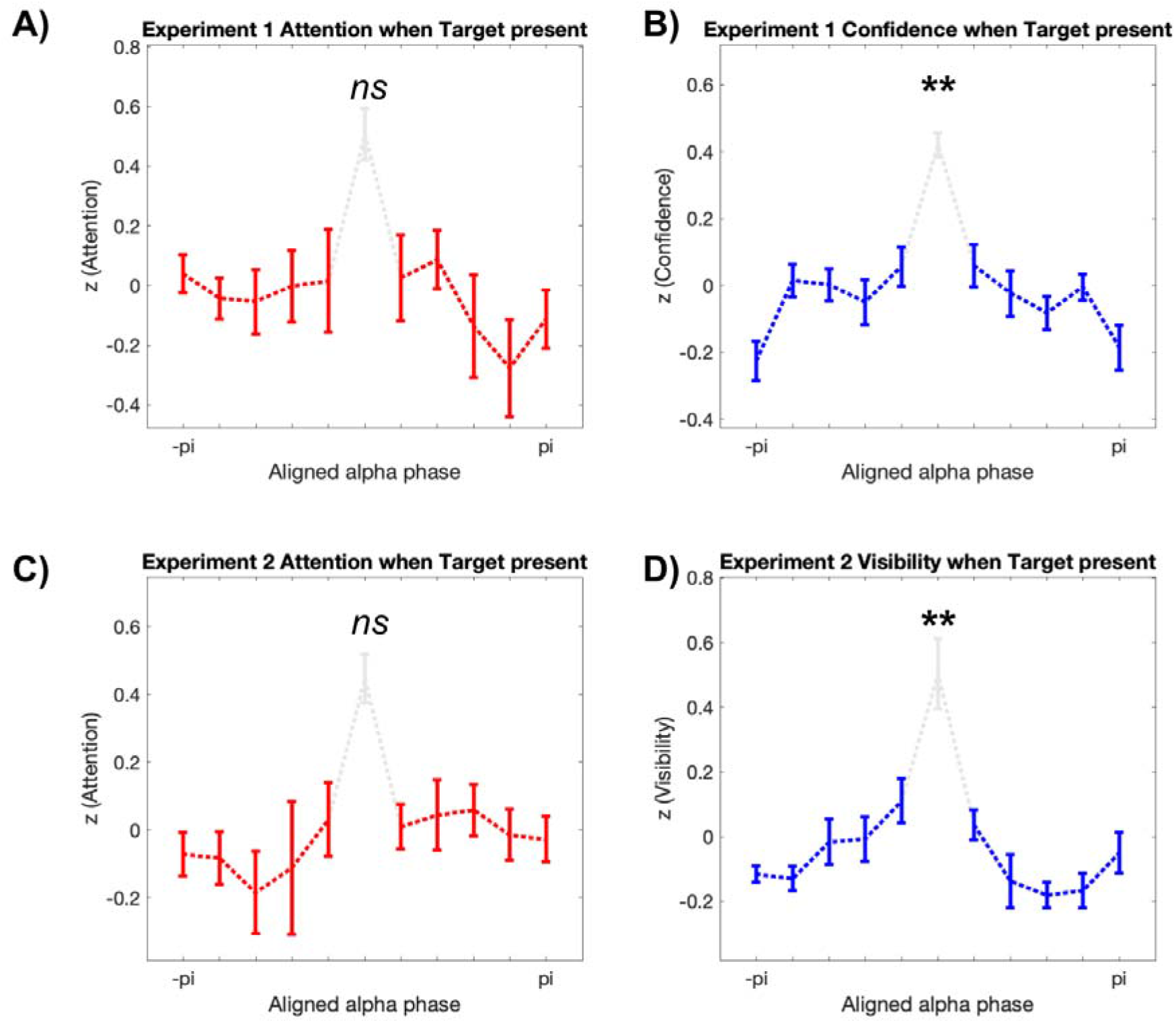
The relationship between preparatory alpha phase at Oz, attention, confidence, and visibility, on target-present trials. A, B) In Experiment 1, only decision confidence varied with alpha phase (Attention: F(4,44) = 1.43, p=.24; Confidence: F(4,44) = 4.24, p = .005). C, D) In Experiment 2, only target visibility varied with prestimulus alpha phase (Attention: F(4,32) = 0.26, p = .9; Visibility: F(4,32) = 4.29, p = .007). ns= not significant, ** p < .01.

